# Genomic and functional characterization of compost-derived bacteria identifies a candidate bioinoculant that influences tomato physiology and leaf metabolism

**DOI:** 10.64898/2026.10.28.762440

**Authors:** Rosalba Cipriani, Susan R. Strickler, Nadine De Biasio, Clarissa Laura Angiolin, Giulia Ghirardello, Werner Zanardi, Tiziano Bonato, Barbara Baldan, Sebastiano Nigris

## Abstract

- Soil biodiversity is increasingly threatened by anthropogenic activities, with intensive agriculture having a particularly detrimental impact on soil microbial communities and potentially disrupting plant-microbe interactions that contribute to plant growth and resilience to environmental stresses. Biofertilizers based on Plant Growth-Promoting Rhizobacteria (PGPR) and organic amendments represent promising strategies to restore microbial diversity and promote crop health. Although compost is widely used as an organic fertilizer, its potential as a reservoir of culturable beneficial bacteria for the development of bioinoculants remains largely unexplored.
- In this work, we isolated and identified six bacterial strains from commercial compost derived from municipal organic waste and screened them for plant growth-promoting traits. Two representative isolates, *Kocuria rhizophila* and *Glutamicibacter arilaitensis*, were subjected to whole-genome sequencing and comparative analyses to investigate their taxonomy, functional potential, rhizosphere competence, secondary metabolism, plastic degradation associated features, and predicted human pathogenicity. *K. rhizophila* was subsequently tested as an inoculant on tomato plants grown in soil, compost-amended soil, or sterile compost-amended soil.
- Five of the six isolates exhibited many of the plant growth-promoting traits tested. Genomic analyses of the two strains supported several of the observed phenotypes and revealed functions potentially associated with nutrient metabolism, plant-root interactions, environmental adaptation, and degradation of plastic polymers. Neither strain was predicted to be pathogenic to humans, indicating a potentially low safety risk for their application as bioinoculants.
- Upon inoculation with *K. rhizophila*, plant responses were influenced primarily by substrate type, although the bacterium produced specific, substrate-dependent effects at 33 days post inoculation. Inoculation resulted in a trend towards increased dry biomass in plants grown in sterile compost and influenced leaf metabolic profiles at 33 days. These results support the potential of *K. rhizophila* as a compost-derived bioinoculant and indicate that enriching compost with beneficial strains isolated from the same matrix could be a useful approach to complement its effects on plant growth.

## 1. Introduction

The long-standing relationship between soil and human society has shaped the development of civilizations, with crop cultivation and food production making soil an indispensable resource for agriculture. In 2023, approximately 38.1% of the total land area of the European Union was used for agriculture (Eurostat, 2026). Although our dependence on soil-water systems is essential for human survival, agricultural practices can also have substantial negative impacts on natural ecosystems. Soil is the most biodiverse habitat on Earth, and it is estimated that 59% of terrestrial life forms are directly linked with soils during their life cycle, making the soil one of the Earth’s most important reservoirs of biodiversity (Anthony *et al*., 2023).

Since the Green Revolution of the 1960s, the intensive use of chemical fertilizers, pesticides, monocultures, and heavy machinery has contributed to meeting the growing global demand for food, but has also had serious consequences for soil biodiversity. Although awareness of the impact of intensive agricultural practices on soil microbial communities has increased over the past 15 years, soil conservation remains insufficient to ensure the long-term maintenance of soil biodiversity, despite the objectives established by the European and global Sustainable Development Agendas (Zeiss *et al*., 2022). Soil biodiversity is closely linked to plant health and productivity, as plants establish numerous interactions with microorganisms that contribute to nutrient acquisition and protection against biotic and abiotic stresses. Greater soil microbial diversity provides plants with access to a wider range of beneficial microorganisms and functions, increasing the possibility of positive plant-microbe interactions (Saleem *et al*., 2019). These interactions occur particularly in the rhizosphere, where microorganisms exchange molecules with plant roots and establish diverse relationships with plants. Well-characterized examples include arbuscular mycorrhizal symbiosis (Ho-Plágaro and García-Garrido, 2022), nitrogen fixation by rhizobia in legumes (Yang *et al*., 2022), and interactions with Plant Growth-Promoting Rhizobacteria (PGPR). PGPR can colonize the rhizosphere and, in some cases, the internal root tissues (endosphere) (Santoyo *et al*., 2021). Their beneficial effects include phytostimulation through the modulation of plant hormones, increased nutrient uptake for plants, and biocontrol through the secretion of antimicrobial compounds or the formation of biofilms (Pathak *et al*., 2026). The ongoing loss of soil microbial biodiversity may therefore reduce the pool of microorganisms capable of forming beneficial interactions with plants (Bloom *et al*., 2025).

In addition to established strategies for preserving soil biodiversity, such as crop rotation and minimal soil tillage, the use of microbial and organic fertilizers represents a promising approach towards more sustainable agricultural practices.

Among these, biofertilizers encompass a wide range of living or dormant microorganisms that provide various benefits to plants, depending on their taxonomic identity and functional profiles. These include microorganisms that solubilize nutrients and PGPR that additionally produce phytohormones and protect plants against soil-borne pathogens (Kumar *et al*., 2022).

Among organic fertilizers, compost is a particularly promising strategy because of its role in both soil improvement and the circular economy. Composting is an aerobic process in which microorganisms decompose organic materials under controlled conditions. An important advantage of compost production is its role in the circular economy, as organic waste (biowaste) can be transformed into a valuable soil amendment while generating products such as water, carbon dioxide, heat, and biogas, which can potentially be recovered for energy production or other applications (Oshins *et al*., 2022). Compost has traditionally been used as a soil amendment to supply organic matter and nutrients, and its use has increased in recent years, partly in response to the growing concerns surrounding the sustainability and the recent price fluctuations of chemical fertilizers (Vos *et al*., 2025). Moreover, compost application can improve soil carbon status and nutrient availability, thereby supporting plant growth and productivity and promoting the proliferation of soil microbial communities (Liu *et al*., 2022). Compost therefore represents a promising tool for maintaining and enhancing soil microbial biodiversity and, consequently, for supporting more sustainable agricultural systems (Heisey *et al*., 2022).

While the beneficial effects of compost application on soil properties, plant growth, and microbial communities are increasingly recognized (Diacono & Montemurro, 2010; Liu *et al*., 2022), comparatively little is known about the potential of compost itself as a source of bacteria with plant growth-promoting properties. Previous research has shown that commercial compost derived from municipal organic waste can be a valuable reservoir of beneficial bacteria displaying multiple plant growth-promoting traits, and that compost application can enhance the growth of grapevine under in vitro conditions (Tondello *et al*., 2022). Building on these findings, in this study we investigated the potential of compost as a source of microbial fertilizers by isolating and biochemically characterizing different bacterial strains. This characterization was complemented by in-depth genomic analyses of two representative strains to investigate their potential as PGPR and identify genomic features potentially associated with adaptation to plant-root interactions. Furthermore, using tomato plants (*Solanum lycopersicum* cv. Micro-Tom), we assessed the effects of inoculating compost and non-compost-amended substrates with a compost-derived bacterium, to evaluate whether bacterial enrichment of compost could influence plant physiology and support the development of more tailored strategies for its agricultural application.

## 2. Materials and methods

### 2.1 Compost

The compost used in this study was produced by S.E.S.A. S.p.A. (Este, Italy) from the organic fraction of municipal waste. Six compost batches were used: M33, M38, M39, M40, M41, and M42. Their physicochemical properties were determined according to UNI standard methods and were compliant with relevant EU directives and consistent with the company’s quality standards (Table S1).

### 2.2 Compost CFU analysis

A colony-forming unit (CFU) analysis was performed to estimate the number of viable bacteria present in the compost. Briefly, 10 g of each of the five compost batches (M38, M39, M40, M41, and M42) was suspended in 50 mL of a sterile MgCl_2_ solution and incubated under agitation on an orbital shaker for 1 h. Serial dilutions were then prepared using the sterile MgCl_2_ solution to obtain suspensions with decreasing bacterial concentrations (1:10, 1:100, 1:1,000, and 1:10,000). Subsequently, 500 μL of the two highest dilutions were plated in triplicate using the spread-plate method on Luria-Bertani (LB) ½ medium and incubated at 28 °C for 4 days. After incubation, colonies were counted, and the counts were multiplied by the corresponding dilution factors to calculate the CFU/g of compost. This procedure was used to assess the viable bacterial load in wet compost. To obtain the corresponding values for dry compost, the moisture content of each batch, as determined by the company (Table S2), was subtracted from the wet compost values.

### 2.3 Isolation and molecular identification of compost-derived bacterial strains

The plates obtained from the CFU analysis were examined for colonies displaying distinct morphological characteristics. Selected colonies were purified by streak plating on LB ½ medium and incubated at 28 °C for 2 days. A small amount of each purified culture was then transferred using a sterile inoculating loop into stock vials containing liquid PCA supplemented with 30% glycerol. The bacterial strains were stored at −80 °C until further analysis.

Genomic DNA was extracted from fresh bacterial cultures using an alkaline lysis method adapted from Zhang *et al*. (2021). Briefly, 20 μL of alkaline lysis buffer (25 mM NaOH and 0.2 mM Na_2_-EDTA, pH 12) was added to a sterile microcentrifuge tube. A single bacterial colony was collected from the culture plate using a sterile pipette tip and suspended in the lysis buffer. The suspension was incubated at 95 °C for 30 min and subsequently cooled to room temperature for a few minutes before being briefly centrifuged. Then, 20 μL of neutralization buffer (40 mM Tris-HCl, pH 7.5) was added, followed by a brief centrifugation.

The 16S rRNA gene was amplified by polymerase chain reaction (PCR) using the universal primers 27F (5′-AGAGTTTGATCCTGGCTCAG-3′) and 1492R (5′-GGTTACCTTGTTACGACTT-3′), and PCR amplification was subsequently verified by electrophoresis on a 1.2% agarose gel (Agarose LE, EuroClone). Selected PCR products were purified using the Exo-SAP method to remove residual primers and unincorporated dNTPs prior to sequencing. For each reaction, 5 μL of PCR product was mixed with 1 μL of Exonuclease I (Exo) and 1 μL of Shrimp Alkaline Phosphatase (SAP) in a microcentrifuge tube. The reaction was incubated at 37 °C for 5 min, followed by incubation at 80 °C for 10 min to inactivate the enzymes.

Purified PCR products were submitted to Eurofins (Germany) for Sanger sequencing. The resulting chromatograms were manually inspected for sequence quality using FinchTV v.1.4.0. Overlapping sequence reads were assembled into a single continuous sequence for each isolate. The resulting sequences were compared against the NCBI database using BLASTn (https://blast.ncbi.nlm.nih.gov/Blast.cgi). For each isolate, the three closest matches were identified based on similarity score and percentage identity, considering matches with >98% identity.

### 2.4 Bacterial growth curves

One bacterial strain representing each identified species was selected for subsequent experiments. Growth curves were generated to establish the relationship between optical density and viable bacterial concentration under the experimental conditions. OD_600_ was measured using a UV-Vis spectrophotometer (Cary 60 UV-Vis, Agilent), and viable cell counts were determined by plate counting. Strains were recovered from −80 °C stocks and inoculated into 5 mL of liquid PCA ½ medium, followed by overnight incubation at 28 °C under agitation. Subsequently, 100 μL of each pre-inoculum was transferred into 10 mL of liquid PCA ½ medium. OD_600_ was measured at 0, 4, 8, 24, and 28 h of incubation at 28 °C under agitation.

At each time point, 0.5 mL of culture was collected for viable cell enumeration. Samples were directly plated and serially diluted in liquid PCA ½ medium (1:2, 1:100, 1:1,000, 1:10,000, and 1:100,000), with 100 μL of each dilution plated on PCA agar using the spread-plate method. Plates were incubated at 28 °C for 24 h, after which colonies were counted and CFU/mL was calculated according to the corresponding dilution factor.

The growth curves were generated by combining OD_600_ and viable cell count data. The linear portion of each curve was identified by visual inspection, and a linear regression equation was calculated for this range. These equations were subsequently used to estimate log_10_(CFU/mL) from OD_600_ measurements in further experiments.

### 2.5 Biochemical assays for plant growth-promoting activities

Selected bacterial strains were evaluated for their potential plant growth-promoting (PGP) activities using qualitative and quantitative biochemical assays for phosphate solubilization, siderophore production, and indole-3-acetic acid (IAA) production.

#### 2.5.1 Bacterial cultures

For each assay, bacterial cultures were prepared by collecting the strains from −80 °C stocks using a sterile inoculation loop and inoculating them into 5 mL of liquid PCA ½ medium. The pre-inocula were incubated overnight at 28 °C under agitation on an orbital shaker. Subsequently, 500 μL of each pre-inoculum was transferred into 10 mL of liquid PCA ½ medium and incubated at 28 °C under agitation. OD_600_ was periodically measured using a UV-Vis spectrophotometer (Cary 60 UV-Vis, Agilent) until the desired bacterial concentration was reached.

#### 2.5.2 Phosphate solubilization

The phosphate-solubilizing ability of the bacterial strains was qualitatively assessed on Pikovskaya agar medium supplemented with 1.5% (w/v) agar (Pikovskaya, 1948). The medium was prepared using 10 g/L glucose, 5 g/L tricalcium phosphate [Ca_3_(PO_4_)_2_], 0.5 g/L ammonium sulfate [(NH_4_)_2_SO_4_], 0.2 g/L sodium chloride (NaCl), 0.1 g/L magnesium sulfate (MgSO_4_·7H_2_O), 0.2 g/L potassium chloride (KCl), 0.002 g/L manganese sulfate (MnSO_4_·H_2_O), 0.002 g/L ferrous sulfate (FeSO_4_·7H_2_O), and 0.5 g/L yeast extract. Bacterial cultures were prepared as described in par. 2.5.1. When cultures reached approximately 10^6^ CFU/mL, 50 μL of each bacterial suspension was spot-inoculated onto Pikovskaya agar plates in triplicate. Plates were incubated at 28 °C for 10 days. Phosphate solubilization was assessed by the formation of a transparent halo around the bacterial colonies.

#### 2.5.3 Siderophore production

Siderophore production was qualitatively assessed using the Blue Agar Chrome Azurol S (CAS) assay according to Louden *et al*. (2011).

Bacterial cultures were prepared as described in par. 2.5.1. When cultures reached approximately 10^6^ CFU/mL, bacterial suspensions were streaked onto CAS agar plates using a sterile inoculation loop. Plates were incubated at 28 °C for 6 days. Siderophore production was assessed based on the formation of an orange/yellow halo around the bacterial growth.

#### 2.5.4 IAA production

IAA production was quantitatively assessed using a colorimetric assay based on the Salkowski reagent method (Gang *et al*., 2019). The Salkowski reagent consisted of 0.5 M ferric chloride (FeCl_3_) and 35% perchloric acid (HClO_4_).

An IAA standard curve (1, 10, 50, and 100 μg/mL) was prepared from an 875.9 μg/mL IAA stock solution in liquid PCA. For each concentration, 1 mL of standard solution was mixed with 1 mL of Salkowski reagent, incubated at 30 °C in the dark for 30 min, and measured at 536 nm using a UV-Vis spectrophotometer (Cary 60 UV-Vis, Agilent). A blank containing liquid PCA and Salkowski reagent was prepared under the same conditions. The resulting standard curve was used to calculate IAA concentrations in the bacterial samples (Table S3).

For the assay, bacterial strains were grown overnight in 5 mL of liquid PCA ½ at 28 °C under agitation. Then, 100 μL of each pre-inoculum was transferred into 10 mL of liquid PCA supplemented with 0.15% (w/v) L-tryptophan and incubated at 28 °C under agitation for 3 days in the dark. When cultures reached at least 10^5^ CFU/mL, as estimated from OD_600_ measurements, 1.5 mL of culture was centrifuged at 16,278 g for 5 min. Subsequently, 1 mL of supernatant was mixed with 1 mL of Salkowski reagent and incubated at 30 °C in the dark for 30 min. Absorbance was then measured at 536 nm, and IAA concentrations were calculated from the standard curve.

### 2.6 Genomic DNA extraction and sequencing of *Kocuria rhizophila* and *Glutamicibacter arilaitensis*

Frozen stocks of *Kocuria rhizophila* and *Glutamicibacter arilaitensis* maintained at −80°C were separately inoculated into 10 mL of LB broth and incubated overnight at 28°C with orbital shaking at 120 rpm. Cells were harvested by centrifugation at 3,400 rpm for 10 min, and the resulting pellets were used for genomic DNA extraction with the DNeasy PowerSoil Kit (QIAGEN, Hilden, Germany), following the manufacturer’s instructions. DNA concentration and purity were estimated using a NanoPhotometer N60 spectrophotometer (Implen, Munich, Germany), with purity evaluated from the A260/A280 and A260/ A230 absorbance ratios.

The genomic DNA samples were submitted to Novogene GmbH (Munich, Germany) for quality control, library preparation and whole-genome sequencing. Continuous long-read (CLR) libraries were prepared and sequenced using the Pacific Biosciences Sequel II platform in CLR mode, targeting 1 Gb of raw sequencing data per sample. Raw-read quality control and read-statistics generation were performed by the sequencing provider.

### 2.7 Genome assembly, annotation and visualization

PacBio sequencing data were obtained as subread BAM files, and raw-read quality was assessed using the Quality Control module of SequelTools v1.3.4 (Hufnagel *et al*., 2020). Circular consensus sequences (CCS) were generated from the subreads using the PacBio ccs tool v8.0.1, implemented in SMRT Link v13.1 (Pacific Biosciences, 2024); the resulting CCS BAM files were converted to compressed FASTQ format using bam2fastq v1.1.0 (Dexheimer, 2010). The consensus reads were independently assembled for *Glutamicibacter arilaitensis* and *Kocuria rhizophila* using hifiasm v0.19.8 (Cheng *et al*., 2021) with default parameters. Primary contigs were extracted and converted to FASTA format. Assembly contiguity and sequence characteristics were evaluated in reference-free mode using QUAST v5.2.0 (Gurevich *et al*., 2013), considering total assembly length, number of contigs, largest-contig length, GC content, N50, L50 and the number of ambiguous bases. Genome completeness was assessed with BUSCO v5.8.2 (Manni *et al*., 2021) in genome mode, using automatic selection of the most appropriate prokaryotic lineage dataset (--auto-lineage-prok). For sequence polishing, the original PacBio subreads were aligned against the corresponding assemblies using pbmm2 v1.17.0 (Pacific Biosciences, 2023) with the SUBREAD preset. The sorted alignments were then used for consensus polishing with gcpp v2.0.2 (Pacific Biosciences, 2020) and the resulting polished assemblies were used in all subsequent analyses.

The taxonomic identity of the two strains was initially assessed by comparing their polished genome assemblies against the NCBI Reference Prokaryotic Genomes database (ref_prok_rep_genomes) using BLASTn. Based on these results, genomes of closely related strains were retrieved for subsequent comparative analyses. Species-level identification was then evaluated using average nucleotide identity (ANI), calculated with FastANI v1.34 (Jain *et al*., 2018). Whole-genome comparisons and ANI supported the assignment of the *Glutamicibacter* isolate to *G. arilaitensis*. The sequenced isolates are therefore referred to hereafter as *G. arilaitensis* M33 and *K. rhizophila* M33. Pairwise comparisons were performed between each M33 genome and the selected reference genomes downloaded from NCBI, using an ANI threshold of 95-96% as the conventional boundary for membership in the same species (Richter and Rosselló-Móra, 2009). The pairwise ANI values were represented as heatmaps, with genomes ordered by average-linkage hierarchical clustering based on ANI-derived distances.

The polished assemblies were structurally annotated using Prokka v1.14.6 (Seemann, 2014), specifying Bacteria as the taxonomic kingdom. Predicted protein sequences were subsequently functionally annotated using eggNOG-mapper v3.0.0-beta6 (Cantalapiedra *et al*., 2021; Hernández-Plaza *et al*., 2026), and the resulting annotations were used to assign predicted functions and Clusters of Orthologous Groups (COG) functional categories.

Putative plasmid sequences were identified from the polished genome assemblies using the *mob_recon* module of MOB-suite v3.1.9 (Robertson and Nash, 2018) with default parameters. Putative plasmid contigs were extracted and annotated separately using Prokka, and their similarity to previously characterized plasmids was investigated through BLASTn searches against the NCBI nucleotide database. A circular representation of the putative plasmid was generated using Proksee (Grant *et al*., 2023) and the Prokka-predicted coding sequences. Potential antimicrobial-resistance determinants were screened using the CARD Resistance Gene Identifier implemented in Proksee, while proteins associated with mobile genetic elements were screened using mobileOG-db with an E-value threshold of 1×10^−20^.

### 2.8 Comparative genomic analysis and in silico functional characterization

Comparative orthology analyses were performed separately for each genus using OrthoFinder v2.5.5 (Emms and Kelly, 2019). Protein sequences predicted by Prokka for the M33 strains and the selected reference genomes were subjected to an all-versus-all sequence similarity analysis, followed by orthogroup assignment and inference of gene trees and species trees. The rooted species trees generated by OrthoFinder were visualized and graphically annotated using the Interactive Tree of Life (iTOL, https://itol.embl.de/).

Candidate M33-specific genes were identified from the OrthoFinder results by comparing each M33 strain with its ten closest selected genomes. *Glutamicibacter arilaitensis* M33 was compared with *G. arilaitensis* ET23-04, LJH19, Re117, JB182, and FME22; *G. mysorens* NBNZ 009 and DSM 12798; and *G. nicotianae* AT6, OTC-16, and DSM 20579. *Kocuria rhizophila* M33 was compared with *K. rhizophila* UNH1, PTI10, Noodlococcus, K12-1, 01AKB, 0819A, KH_056, NBC_01227, NBC_00874, and NBC_00863. Candidate M33-specific genes comprised genes assigned to orthogroups present in the corresponding M33 genome but absent from all ten comparator genomes, together with M33 genes that were not assigned to an orthogroup by OrthoFinder. The protein sequences encoded by these candidate genes were functionally annotated using eggNOG-mapper v3.0.0-beta6, and the resulting annotations were used to assign predicted functions and Clusters of Orthologous Groups (COG) functional categories.

Circular representations of the bacterial genomes were generated using Proksee. Custom GFF3 tracks derived from the OrthoFinder analysis were added to the chromosomal maps to show the distribution of M33 genes. For *G. arilaitensis* M33, tracks were generated for strains ET23-04, FME22, LJH19 and Re117, whereas *K. rhizophila* M33 was compared with strains NBC_00874, NBC_01227, KH_056 and 0819A.

The functional potential of the M33 strains was investigated using several parallel genome-mining approaches. Plant growth-promoting traits were predicted using PLaBAse v1.02 (Patz *et al*., 2021), with Prokka-predicted proteins analyzed by both relaxed BLASTP and strict BLASTP-HMMER searches. The strict results were used to classify proteins according to PLaBAse categories for direct and indirect plant growth-promoting effects. Biosynthetic gene clusters potentially involved in specialized metabolite production were identified in the two M33 genome assemblies using antiSMASH v8.0.4 (Blin *et al*., 2025). Analyses were conducted using both strict and relaxed detection settings, while the strict results were used for the conservative reporting of predicted clusters, classified by biosynthetic type.

Rhizosphere-competence-related catabolic gene clusters (rCGCs) were investigated using rhizoSMASH v0.4.0 (Li *et al*., 2025) under strict detection settings in the M33 and reference genomes. rCGC types were counted and compiled into strain-by-rCGC matrices, which were visualized as heatmaps. Strains were clustered by complete-linkage hierarchical clustering based on Euclidean distances of rCGC profiles, and rCGCs were ordered by their distribution across the analyzed genomes.

Plastic degradation potential was assessed using the PlasticDB Compare Genomes tool (Gambarini *et al*., 2022). Prokka-predicted proteins from the M33 and reference genomes were compared against PlasticDB proteins by BLASTP, with *Ideonella sakaiensis* included as a reference. Hits were grouped by polymer, and normalized hit values were used for comparative analysis and bubble-plot visualization.

Finally, the M33 genome assemblies were screened for putative virulence-associated sequences by BLASTn against the core dataset (Set A) of the Virulence Factor Database (VFDB; Zhou *et al*., 2025). Matches were evaluated according to nucleotide identity, alignment length, query coverage, and E-value. Potential human pathogenic capacity was further assessed using PathogenFinder2 (Ferrer Florensa *et al*., 2026), which provides computational pathogenicity scores and binary classifications. Both analyses were interpreted as genomic predictions rather than experimental evidence of virulence or pathogenicity.

### 2.9 Experimental setup and plant inoculation with *K. rhizophila* M33

*Solanum lycopersicum* L. cv. Micro-Tom seeds were surface-sterilized in 5% sodium hypochlorite on an orbital shaker for 20 min, followed by four washes with sterile water. Seeds were then imbibed in sterile tap water and left under agitation for 48 h at room temperature before being sown on half-strength Murashige and Skoog (MS) basal medium supplemented with 1.5% agar. Seeds were germinated in vitro at 25°C and 65% relative humidity under a 16 h light/8 h dark photoperiod. After 12 days, seedlings were transferred to 0.5-L plastic pots containing different soil treatments corresponding to six experimental conditions: soil, soil supplemented with 33 g/L compost, soil supplemented with 33 g/L sterile compost, and the corresponding three conditions with *Kocuria rhizophila* M33 inoculation.

The baseline soil consisted in a mixture composed of 45% agricultural field soil, 45% peat, 5% perlite, and 5% sand. Its physicochemical properties were determined according to UNI standard methods (Table S4). Sterile compost was obtained by autoclaving at 121°C for 2 h. Sterility was assessed by suspending 10 g of sterile compost in 50 mL of sterile deionized water and shaking the suspension on an orbital shaker for 1 h. An aliquot of 100 µL was spread onto Plate Count Agar (PCA) and incubated at 28 °C for 4 days. No colonies were detected under these culture conditions, confirming successful sterilization. The compost concentration of 33 g/L was selected based on previous experiments performed by the research group.

*K. rhizophila* M33 was recovered from a −80°C stock and grown overnight on PCA agar at 28°C. Bacteria were then collected from the plates and used to inoculate 5 mL of liquid PCA. This resulting pre-inoculum was incubated overnight at 28°C with agitation and subsequently transferred to 200 mL of liquid PCA for overnight cultivation under the same conditions. Cells were harvested by centrifugation and resuspended in Murashige and Skoog medium to obtain a final concentration of 10^6^ CFU/mL. Plants assigned to the inoculated treatments received 5 mL of bacterial suspension one day after transplantation, with a second inoculation performed 15 days later. Plants were regularly sub-irrigated with tap water.

Twenty biological replicates were established for each treatment. Ten plants per treatment were destructively sampled after 14 days, while the remaining plants were sampled 19 days later, at 33 days of growth and following the second bacterial inoculation. At each sampling point, leaves and roots were collected. Leaf samples from three plants per treatment were collected for metabolic analyses, ground in liquid nitrogen, and stored at −80°C until further analysis.

### 2.10 Physiological measurements and statistical analysis

Leaf and root fresh biomass was measured immediately after sampling. Samples were then dried at 60 °C for seven days and weighed again to determine dry biomass. Water content was calculated as (fresh weight – dry weight)/dry weight. Photosynthetic efficiency was measured at 14 and 33 days using a MultispeQ V2.0 fluorometer without dark acclimation and expressed as the F_v_’/F_m_’ ratio, representing the maximum potential quantum efficiency of photosystem II under light conditions.

At both sampling times, metabolic markers, including photosynthetic pigments, free amino acids, soluble sugars, starch, and phenolic compounds were quantified using the colorimetric “rainbow protocol” (López-Hidalgo *et al*., 2021). Frozen leaf material was lyophilized, and 8 mg of tissue were homogenized prior to extraction. Three biological replicates were analyzed for each treatment, with three technical replicates per biological replicate.

Statistical analyses were conducted in R version 4.6.1 (R Core Team, 2026). Leaf and root biomass, water content and photosynthetic efficiency were compared between *Kocuria rhizophila*-inoculated and non-inoculated plants within each substrate and sampling time using two-sided unpaired Wilcoxon rank-sum tests. Statistical significance was set at p < 0.05. Metabolic profiles obtained from the rainbow protocol were further analyzed separately at 14 and 33 days using principal component analysis (PCA) after standardization of the measured variables. The effects of inoculation and its interaction with substrate were assessed by PERMANOVA based on Euclidean distances with 9,999 permutations restricted within substrate. Results were visualized using boxplots showing individual observations and PCA score plots.

## 3. Results

### 3.1 Culturable bacterial load, strain identification and growth characterization

To estimate the abundance of viable, culturable bacteria in the compost batches, culturable bacterial loads were determined and expressed as log(CFU/g) on a wet-weight basis. The values were then normalized to the dry weight of each batch using the supplier-reported moisture content. For batch M38, the reported value was based on the only plate with a colony count within the quantifiable range. Among the five compost batches, M38 showed the highest culturable bacterial load, reaching 8.9 log (CFU/g) dry compost, followed by M39, M40, M41, and M42 (Table S5). The culturable bacterial loads calculated for wet and dry compost batches are presented in Figure 1A.

**Figure 1.**
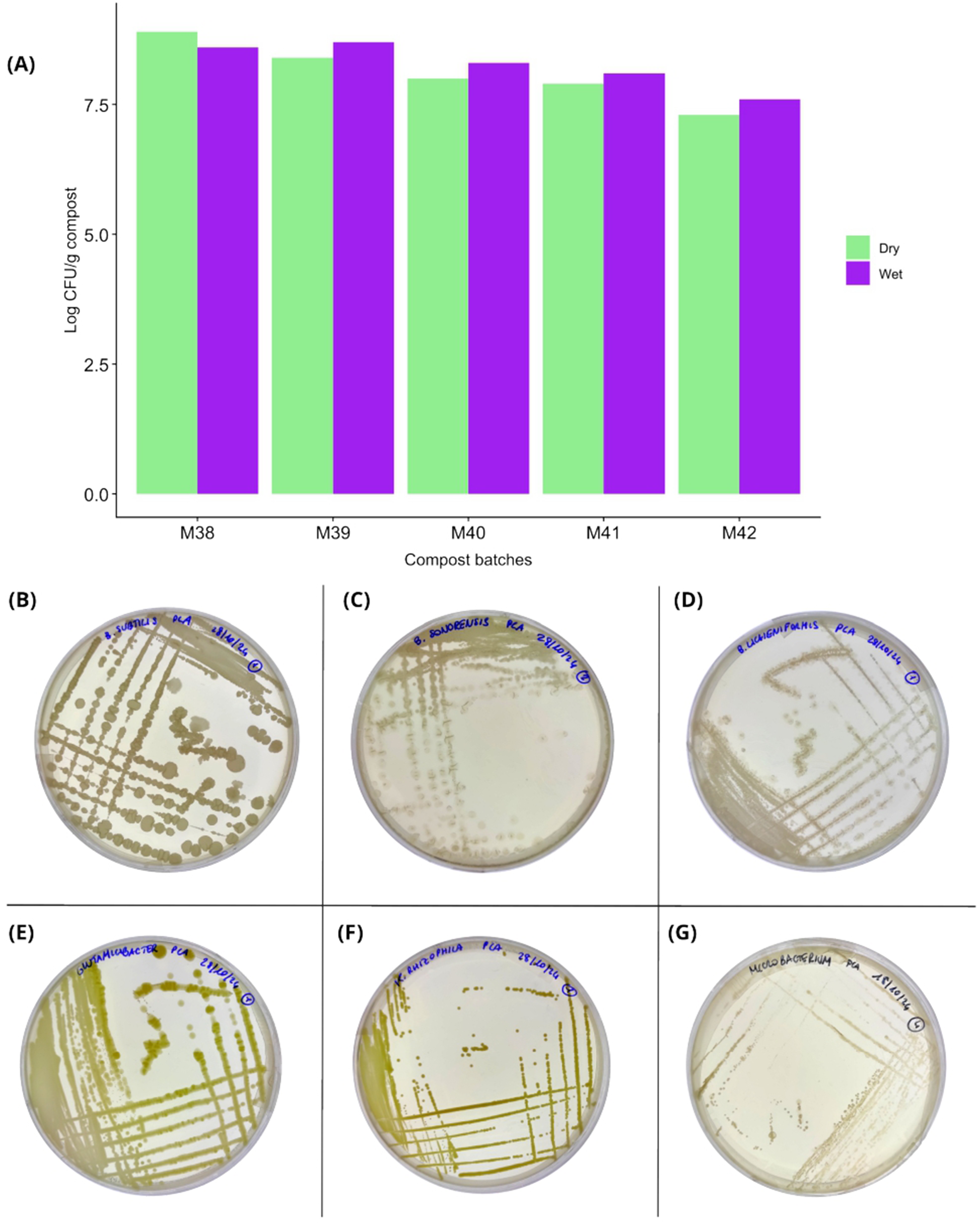
Bacterial load and representative bacterial isolates recovered from compost samples. **(A)** Quantification of culturable bacteria in the different compost batches, expressed as log CFU/g of compost. Bars represent the measured bacterial abundance per gram of wet and dry compost batches. **(B–G)** Representative photographs of agar plates showing the six bacterial isolates recovered from the compost samples. Each panel corresponds to one bacterial isolate: **(B)** *Bacillus subtilis*, **(C)** *Bacillus sonorensis*, **(D)** *Bacillus licheniformis*, (**E**) *Glutamicibacter sp.*, (**F**) *Kocuria rhizophila*, and (**G**) *Microbacterium suwonense*. The plate images illustrate the colony morphology and growth characteristics of the selected isolates.

In parallel with quantification, a collection of culturable bacterial strains was established. Colonies were selected based on morphological characteristics, including shape, color, and elevation, and subsequently purified by subculturing.

Taxonomic identification of the strains in the different samples and the results of the sequence similarity analysis are summarized in Table S6. Most of the isolates belonged to the genus *Bacillus*, with *B. subtilis* representing the most frequently isolated species, followed by *B. licheniformis* and *B. sonorensis*. The remaining isolates were assigned to *Kocuria rhizophila*, *Microbacterium suwonense*, and *Glutamicibacter* sp., for a total of six bacterial taxa (Figure 1B-G). For isolate *M40-4 (1)_1*, sequence-similarity analysis returned only a limited number of matches to *Glutamicibacter*, including *G. arilaitensis* and an unidentified *Glutamicibacter* species. Given the small number of supporting hits, this initial identification of the strain was limited to the genus.

Bacterial growth kinetic characterization showed that during the first 28 h of culture, all six compost-isolated strains showed increased OD_600_ values, although the magnitude differed among isolates (Figure 2, Table S7). At 28 h, *K. rhizophila* and *Glutamicibacter* sp. showed the highest turbidity (OD_600_ 1.98 and 1.81, respectively), whereas *M. suwonense* remained lower (OD_600_ 0.86) (Fig. 2A). CFU counts generally increased over time, with *Glutamicibacter* sp., *K. rhizophila*, *B. licheniformis*, and *B. subtilis* reaching 8.86, 8.54, 8.19, and 8.17 log_10_(CFU/mL), respectively, at 28 h. *B. sonorensis* reached 7.40 log_10_(CFU/mL), while *M. suwonense* peaked at 9.17 log_10_(CFU/mL) at 24 h before declining to 7.32 log_10_(CFU/mL) at 28 h (Fig. 2B).

**Figure 2.**
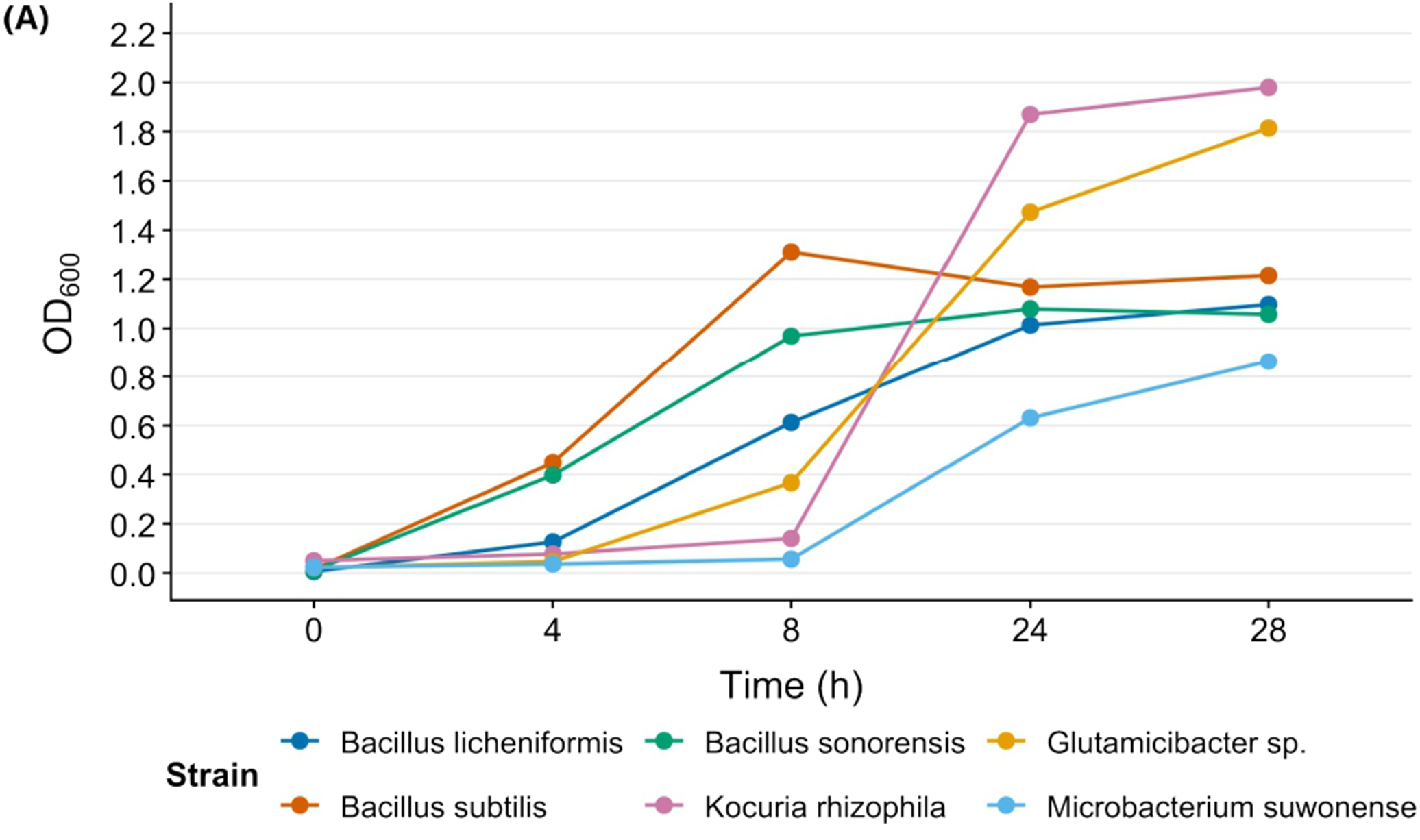
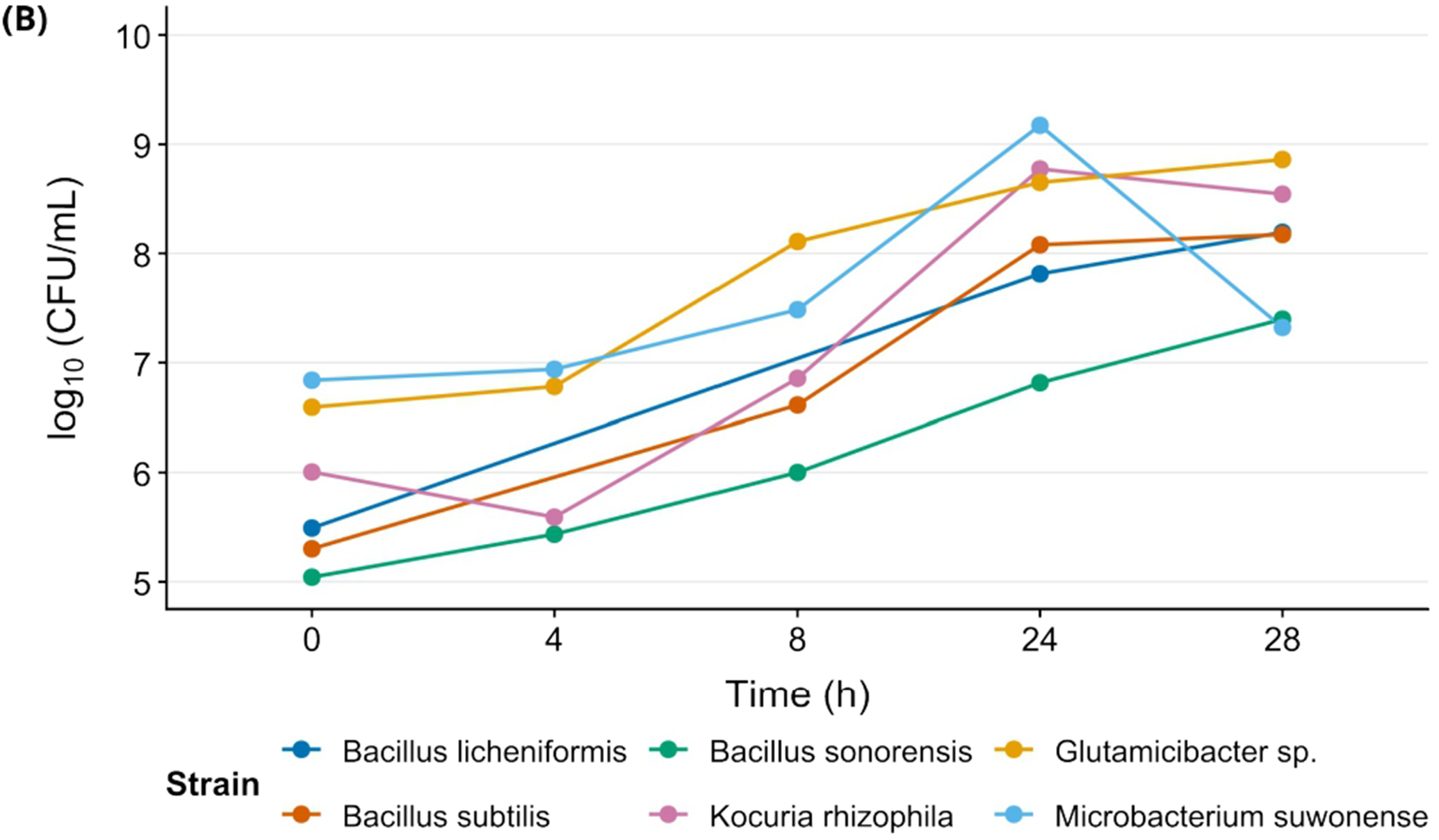
Growth of compost-isolated bacterial strains over 28 h. **(A)** Optical-density growth curves measured at 600 nm (OD_600_). **(B)** Viable-cell growth curves expressed as log_10_(CFU/mL). Curves are shown for *Bacillus licheniformis*, *Bacillus subtilis*, *Bacillus sonorensis*, *Kocuria rhizophila*, *Glutamicibacter* sp., and *Microbacterium suwonense*. Symbols represent measurements at each sampling time; lines connect consecutive time points. CFU measurements were unavailable at some intermediate time points for *B. licheniformis* and *B. subtilis*; therefore, the connecting lines represent trajectories between available observations.

### 3.2 Biochemical characterization of compost-isolated bacteria revealed promising plant growth-promoting traits

With the aim of elucidating the potential of the isolated strains to interact with plants, three biochemical assays were performed; the results are reported in Table 1. Phosphate solubilization was observed in three of the six isolates (Figure S1A-C), with *B. sonorensis* showing the strongest response, followed by *K. rhizophila* and *B. subtilis*. Siderophore production was detected in all isolates except *B. licheniformis* (Fig. S1D-F). The strongest responses were observed for *B. sonorensis* and *Glutamicibacter* sp., whereas *K. rhizophila* showed intermediate activity.

**Table 1.** Plant growth-promoting traits of the selected bacterial isolates. Phosphate solubilization, siderophore production, and indole-3-acetic acid (IAA) production are shown as qualitative responses and indicated as absent (−) or by increasing activity (+ to +++), while IAA concentrations are reported in µg/mL.

| STRAIN | PHOSPHATE SOLUBILIZATION | SIDEROPHORES PRODUCTION | IAA PRODUCTION | IAA (µg/mL) |
| --- | --- | --- | --- | --- |
| *Bacillus licheniformis* | - | - | - | 0 |
| *Bacillus subtilis* | + | + | + | 3.77 |
| *Bacillus sonorensis* | +++ | +++ | + | 2.04 |
| *Kocuria rhizophila* | ++ | ++ | + | 7.36 |
| *Glutamicibacter* sp. | - | +++ | ++ | 34.69 |
| *Microbacterium suwonense* | - | + | +++ | 44.10 |

IAA production was detected in five isolates and ranged from 2.04 to 44.10 µg/mL. *M. suwonense* produced the highest IAA concentration (44.10 µg/mL), followed by *Glutamicibacter* sp. (34.79 µg/mL). Lower concentrations were recorded for *K. rhizophila* (7.36 µg/mL), *B. subtilis* (3.77 µg/mL), and *B. sonorensis* (2.04 µg/mL). *B. licheniformis* showed none of the evaluated activities and did not produce detectable IAA (Table S8).

### 3.3 General features and phylogenetic analysis of *Glutamicibacter arilaitensis* M33 and *Kocuria rhizophila* M33

The assembly obtained with hifiasm generated two genomes of 3,622,309 bp for *Glutamicibacter* sp. and 2,724,449 bp for *K. rhizophila*. The genetic features and BUSCO completeness are summarized in Table 2. Prokka annotation predicted 2333 CDSs, 9 rRNAs, 53 tRNAs and 1 tmRNA for *K. rhizophila*, and 3312 CDSs, 19 rRNAs, 67 tRNAs and 1 tmRNA for *Glutamicibacter* sp.

**Table 2.** Genome assembly statistics and completeness assessment for *Glutamicibacter* sp. and *Kocuria rhizophila*. Assembly contiguity is reported using N50 and L50 values, together with the number of contigs, GC content, total assembly length, and the percentages of single-copy, duplicated, fragmented, and missing Benchmarking Universal Single-Copy Orthologs (BUSCOs). BUSCO analyses were performed using the Micrococcaceae and *Kocuria* lineage datasets for *G. arilaitensis* and *K. rhizophila*, respectively.

| Features | *Glutamicibacter* sp. | *Kocuria rhizophila* |
| --- | --- | --- |
| **N50** | 3,514,832 | 2,671,791 |
| **L50** | 1 | 1 |
| **N. of contigs** | 3 | 2 |
| **GC content (%)** | 59.22 | 70.61 |
| **Total length (bp)** | 3,622,309 | 2,724,449 |
| **BUSCO Single copy genes (%)** | 98.6 (database *micrococcaceae*) | 99.2 (database *kocuria*) |
| **BUSCO Duplicated genes (%)** | 0.3 | 0 |
| **BUSCO Fragmented genes (%)** | 0.5 | 0.2 |
| **BUSCO Missing genes (%)** | 0.5 | 0.6 |

BLAST searches of both genome assemblies identified the most closely related reference strains, and BLAST results also supported the identification of the *Glutamicibacter* isolate as *Glutamicibacter arilaitensis*. OrthoFinder was subsequently used to compare the gene content of each isolate with that of related genomes (22 *Glutamicibacter* genomes and 28 *Kocuria* genomes in total) and to assess their phylogenetic relationships based on shared orthogroups. Both isolates were named M33 after the compost batch from which they were isolated.

For *Glutamicibacter* sp., 73,766 genes were assigned to 5,715 orthogroups, of which 1,910 were shared across all 22 compared genomes, and 1,585 were single-copy orthogroups. In *G. arilaitensis* M33, 3,271 of the 3,312 predicted genes (98.8%) were assigned to 3,124 orthogroups, whereas 41 genes remained unassigned (Table S9). Pairwise average nucleotide identity values were calculated using FastANI and are presented as a heatmap. The orthogroup-based phylogeny placed M33 within the *G. arilaitensis* cluster and close to strain ET23-04, further supporting its species-level identification (Figure 3).

**Figure 3.**
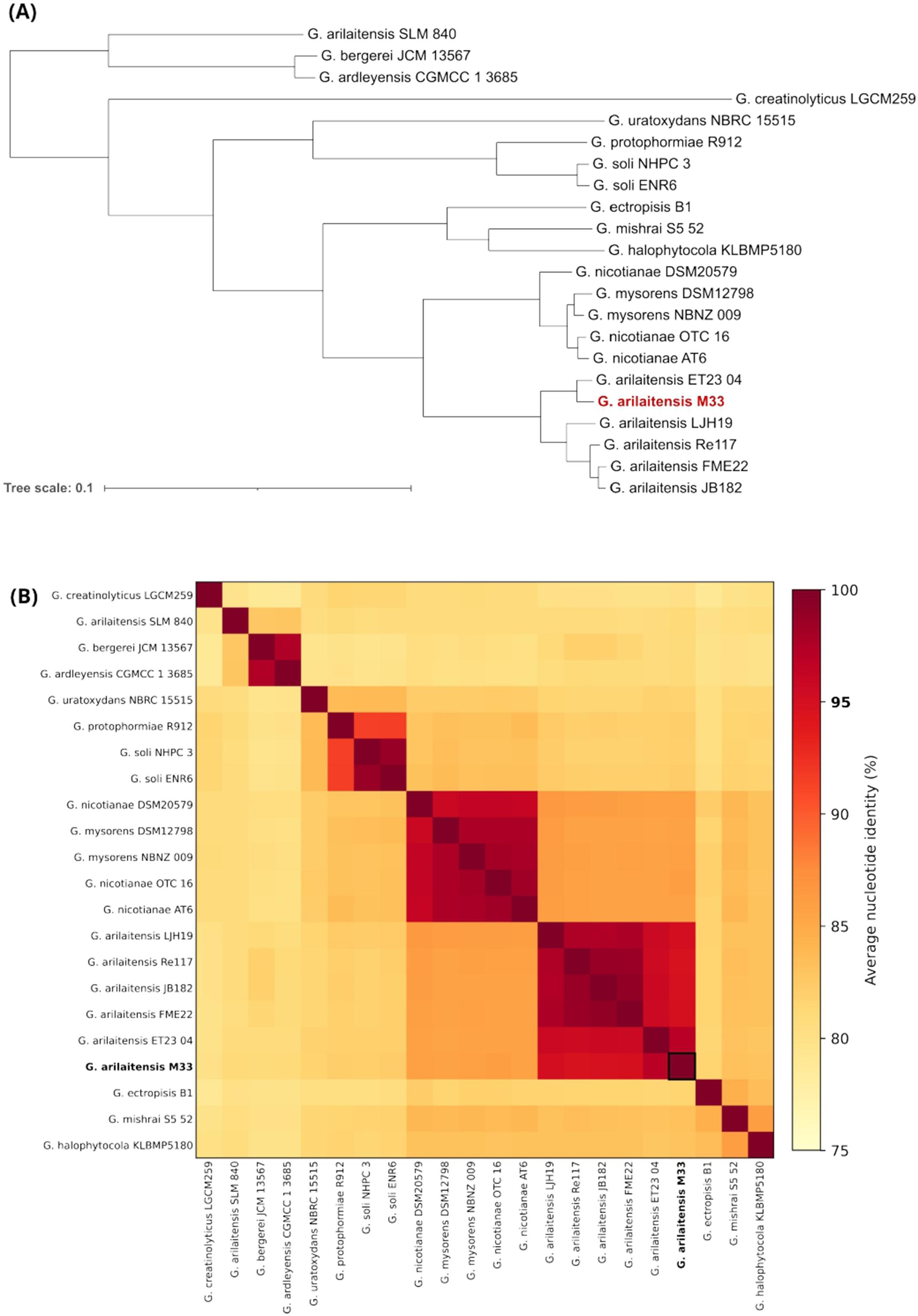
Phylogenetic tree and average nucleotide identity of *Glutamicibacter arilaitensis* M33. **(A)** Species tree showing the relationships among 22 *Glutamicibacter* genomes based on shared orthogroups identified using OrthoFinder. **(B)** Pairwise average nucleotide identity (ANI) heatmap of the *Glutamicibacter* genomes. Colours indicate ANI values ranging from 75% to 100%, with the bold 95% tick marking the commonly used species-level threshold.

For *K. rhizophila* M33, 72,402 genes from 28 genomes were assigned to 4,944 orthogroups, including 1,070 single-copy orthogroups. Of the 2,333 genes predicted in M33, 2,288 (98.1%) were assigned to 2,245 orthogroups, while 45 remained unassigned (Table S10). Average nucleotide identity was calculated and reported as well. The phylogenetic tree placed M33 within the *K. rhizophila* cluster, near strains NBC 00863 and NBC 00874 (Figure 4).

**Figure 4.**
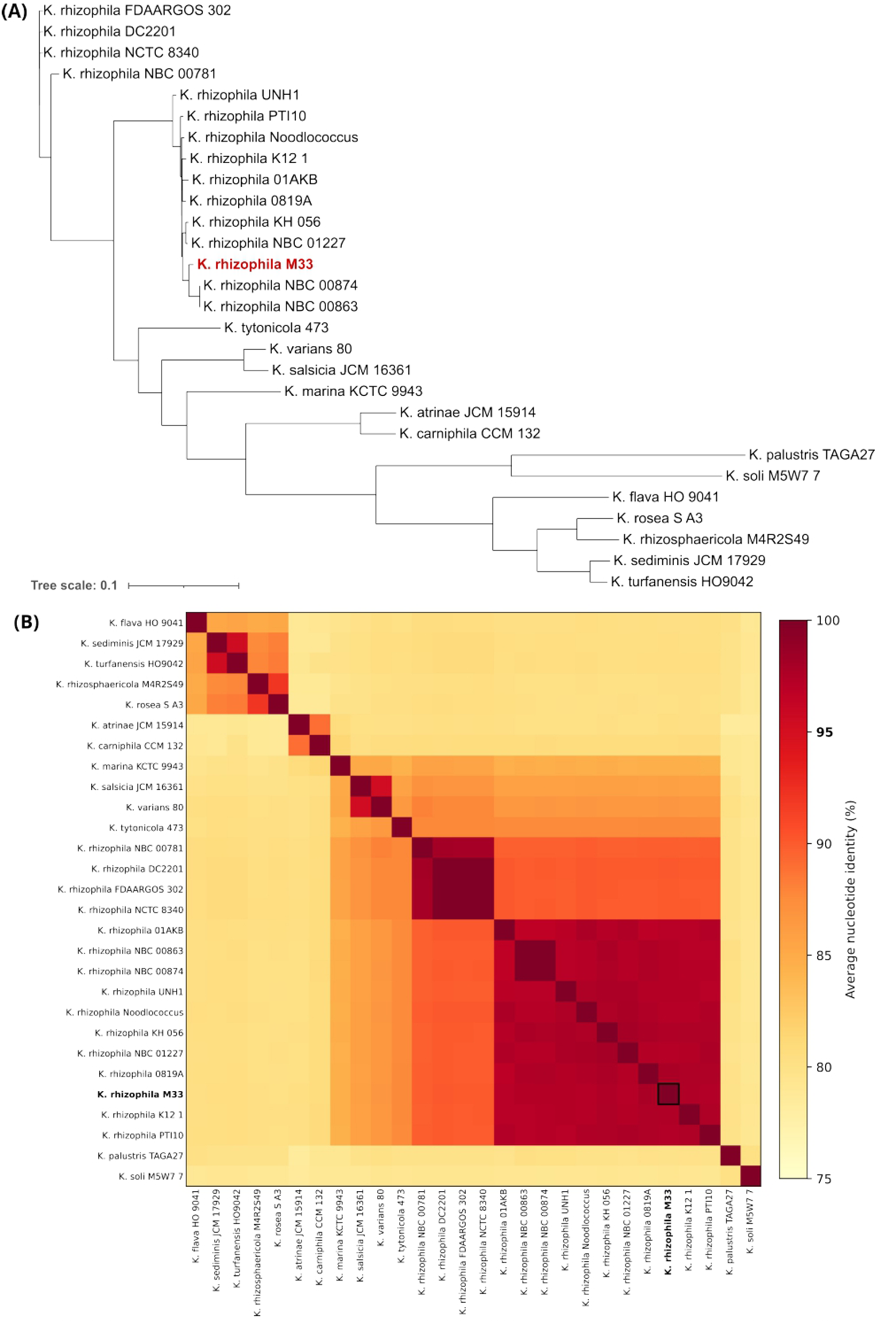
Phylogenetic tree and average nucleotide identity of *Kocuria rhizophila* M33. **(A)** Species tree showing the relationships among 28 *Kocuria* genomes based on shared orthogroups identified using OrthoFinder. **(B)** Pairwise average nucleotide identity (ANI) heatmap of the *Kocuria* genomes. Colours indicate ANI values ranging from 75% to 100%, with the bold 95% tick marking the commonly used species-level threshold.

### 3.4 Plasmid detection and analysis

MOB-suite analysis of the *Glutamicibacter* genome assembly identified one putative plasmid contig, with a length of 98,165 bp and a GC content of 56.91%. The contig was assigned to MOB cluster AC261, with the closest reference sequence belonging to *Glutamicibacter nicotianae*. No replicon, relaxase, mating-pair formation system, or oriT type was assigned, and plasmid mobility could not be predicted.

BLASTn analysis showed that the putative plasmid was most similar to plasmid pGnpA106 of *G. nicotianae* strain OTC-16 (CP033082.2), with 99.17% nucleotide identity across 67% of the query sequence. The second-best plasmid match was the unnamed plasmid of *Glutamicibacter* sp. FR1 (CP178603.1), showing 89.36% identity and 48% query coverage. The remaining plasmid-associated matches covered only 1-4% of the query (Table S11). Thus, the detected contig appears closely related to pGnpA106 but cannot be considered identical because approximately one-third of its sequence was not covered by the alignment.

In contrast, MOB-suite did not identify any putative plasmid contigs in the *K. rhizophila* genome assembly.

The putative plasmid identified in the *Glutamicibacter* genome was visualized and annotated using Proksee (Figure S2), with coding sequences predicted by Prokka and potential antimicrobial-resistance determinants screened against CARD.

Prokka annotation identified 112 coding sequences. 97 (86.6%) encoded hypothetical proteins and 15 (13.4%) were assigned a predicted function. A mercury-resistance region comprising *merR1*, *merA*, and *merB* was identified, suggesting a potential role in the regulation and detoxification of inorganic and organic mercury compounds. The plasmid also contained a group of genes associated with aromatic-compound metabolism.

The CARD analysis returned 77 matches (classified as Loose) associated with 66 distinct resistance ontology entries. The most frequently represented mechanisms were antibiotic inactivation and efflux, followed by target alteration. The matches were associated mainly with β-lactams, including penicillins, cephalosporins, carbapenems, and monobactams, as well as macrolides, tetracyclines, aminoglycosides, fluoroquinolones and phenicols (Table S12).

MobileOG screening (E-value 1E-20) identified six mobile genetic element-associated loci on the putative *Glutamicibacter* plasmid, with sequence identities of 98.8-100%. Three were associated with replication, recombination, and repair functions (*umuD*, *impB*, and *parA*), while three were assigned to the transfer category. However, they do not demonstrate the presence of a complete or functional conjugation system. This interpretation is consistent with the MOB-suite results, which did not assign a mobility class to the plasmid.

### 3.5 Genome-wide orthogroup distribution and functional characterization

With the aim of investigating strain-specific genes, the genomic distribution of M33-specific genes was examined within the phylogenetic clade containing each isolate. For both *G. arilaitensis* M33 and *K. rhizophila* M33, these genes were identified by comparison with ten genomes from the respective M33-containing clade. COG functional categories were assigned to 40 of the 76 candidate M33-specific genes in *G. arilaitensis* and to 25 of the 62 candidate M33-specific genes in *K. rhizophila* and their functional annotations are represented in Figure S3.

Four selected closely related strains were subsequently used for genome-wide visualization. *G. arilaitensis* M33 was compared with strains ET23-04, LJH19, FME22 and Re117 (Figure 5A), whereas *K. rhizophila* M33 was compared with strains KH_056, NBC_00874, 0819A and NBC_01227 (Fig. 6A).

**Figure 5.**
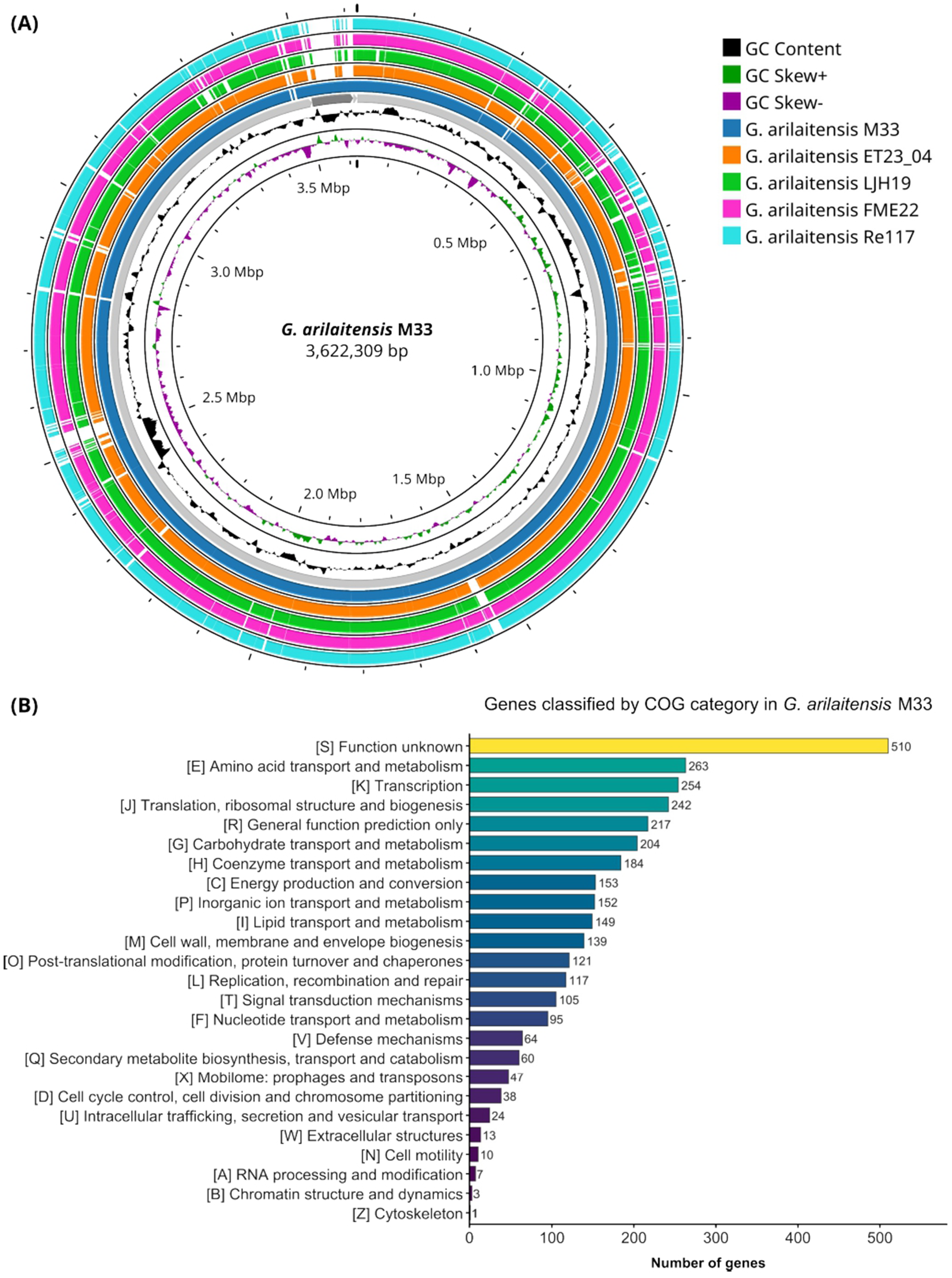
Comparative genomic mapping of *G. arilaitensis* M33 and functional annotation of the M33 genome. **(A)** Proksee circular map of *G. arilaitensis* M33 genome. The two innermost tracks show GC skew and GC content. The subsequent rings, from inner to outer, represent M33 and the four closely related strains ET23-04, LJH19, FME22 and Re117. Tracks represent orthogroup presence obtained with OrthoFinder. **(B)** Distribution of M33 genes among COG functional categories based on eggNOG-mapper annotations. Bars indicate the number of genes assigned to each category and are ordered by gene count. Genes assigned to multiple COG categories were counted once in each corresponding category

**Figure 6.**
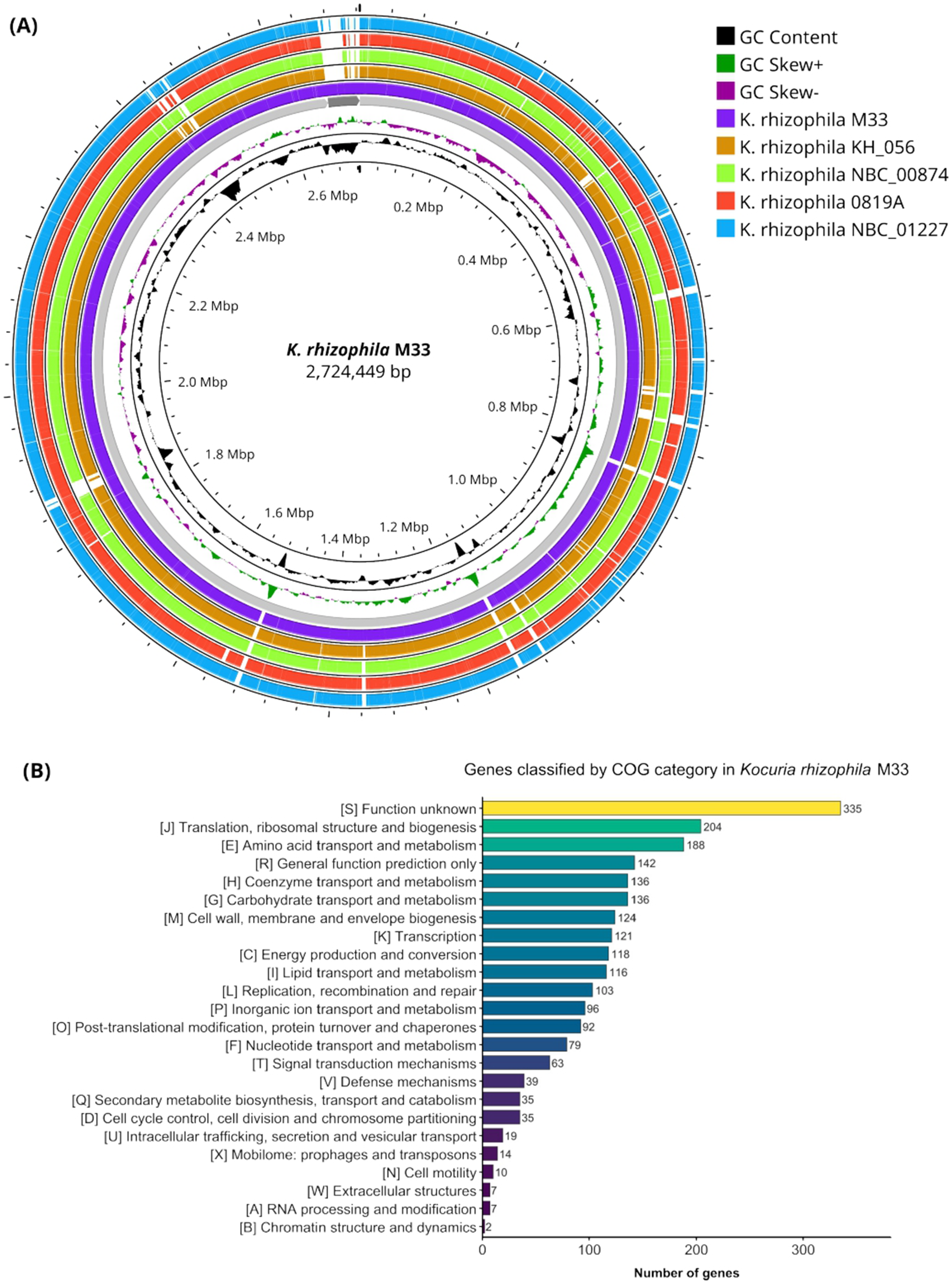
Comparative genomic mapping of *K. rhizophila* M33 and functional annotation of the M33 genome. **(A)** Proksee circular map of *K. rhizophila* M33 genome. The two innermost tracks show GC skew and GC content. The subsequent rings, from inner to outer, represent M33 and the four closely related strains KH_056, NBC_00874, 0819A and NBC_01227. Tracks represent orthogroup presence obtained with OrthoFinder. **(B)** Distribution of M33 genes among COG functional categories based on eggNOG-mapper annotations. Bars indicate the number of genes assigned to each category and are ordered by gene count. Genes assigned to multiple COG categories were counted once in each corresponding category.

Orthogroup distribution was conserved across most regions of both M33 genomes. The comparison tracks also showed some gaps, indicating loci for which corresponding orthologues were absent from one or more of the selected reference genomes. Regions showing gaps across all four reference tracks included M33-specific genes identified through the broader clade-level comparisons.

EggNOG functional characterization of the M33 genomes showed that genes of unknown function constituted the largest category in both strains (Figg. 5B and 6B). Among functionally characterized genes, amino acid transport and metabolism, transcription, general function prediction, and carbohydrate transport and metabolism were highly represented in both genomes. Signal transduction mechanisms constituted an additional prominent category in *G. arilaitensis* M33, whereas translation, ribosomal structure and biogenesis were particularly represented in *K. rhizophila* M33.

### 3.6 Genomic prediction of plant growth-promoting traits

To support and complement the plant growth-promoting phenotypes observed in the biochemical assays, the genomes of *K. rhizophila* M33 and *G. arilaitensis* M33 were screened against PLaBAse using combined BLASTP and HMMER searches. PLaBAse identified 2,371 and 3,489 plant growth-promoting trait annotation occurrences in *K. rhizophila* M33 and *G. arilaitensis* M33, respectively; Figure 7 represents hierarchical annotation frequencies.

**Figure 7.**
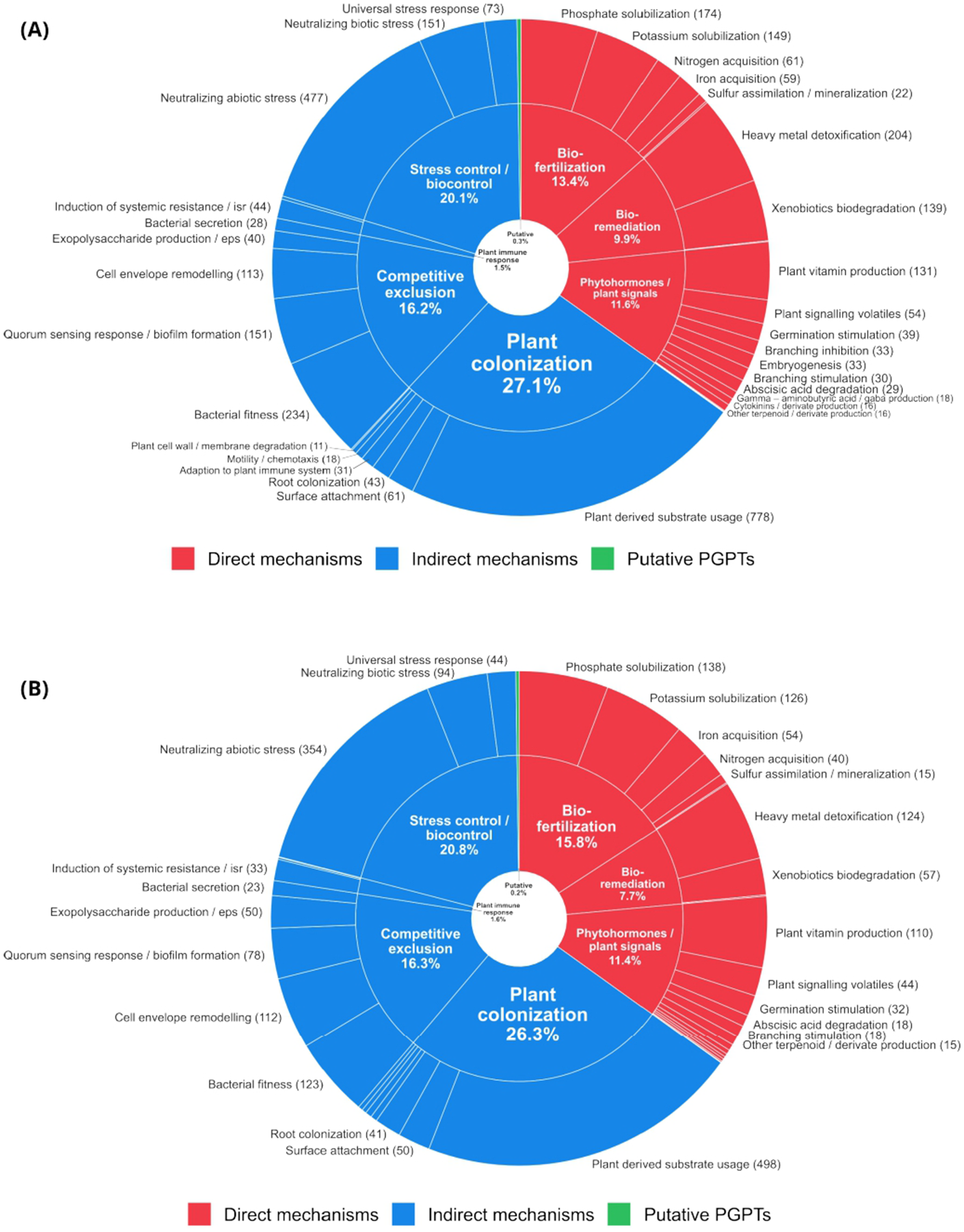
Predicted plant growth-promoting traits in *G. arilaitensis* M33 and *K. rhizophila* M33. Sunburst plots summarizing plant growth-promoting trait (PGPT) annotations identified by PLaBAse using combined BLASTP and HMMER searches in **(A)** *G. arilaitensis* M33 and **(B)** *K. rhizophila* M33. The inner ring shows broad functional categories, with percentages indicating their contribution to the total PLaBAse annotation frequency. The outer ring shows the corresponding specific functions. All functions are represented by sectors, whereas external labels are displayed only for functions with a frequency greater than 10. Colours distinguish direct mechanisms, indirect mechanisms and putative PGPTs.

The analysis conducted for *K. rhizophila* M33 showed that the positive phosphate-solubilization phenotype was supported by 138 assignments, mainly related to organic-acid metabolism, phosphatase activity, and phosphate transport or homeostasis. In contrast, *G. arilaitensis* M33, which was negative in the phenotypic assay, showed 174 phosphate-solubilization assignments, indicating that these functions do not necessarily result in detectable activity under the tested conditions. IAA-related annotations were detected in both genomes (18 in *K. rhizophila* M33 and 29 in *G. arilaitensis* M33), while siderophore-related annotations accounted for 16 and 14 occurrences, respectively, including the presence of desferrioxamine, an important family of hydroxamate-type siderophores.

Further validation was performed through strict antiSMASH analysis for secondary metabolites screening (Table S13). This analysis identified five and six biosynthetic regions in *K. rhizophila* M33 and *G. arilaitensis* M33, respectively. Overall, the PLaBAse and antiSMASH analyses supported the observed IAA and siderophore phenotypes and phosphate solubilization by *K. rhizophila* M33, revealing a genotype-phenotype discrepancy for phosphate solubilization in *G. arilaitensis* M33.

### 3.7 Investigation of rhizosphere-competence-related catabolic gene clusters (rCGCs)

As a further step in the characterization of these isolates, we investigated their predicted root-associated functional potential by analysing root catabolic gene clusters (rCGCs), with the aim of identifying genomic features that could contribute to interactions with plant roots (Figure 8). *G. arilaitensis* M33 (Fig. 8A) showed a relatively rich and diverse rCGC repertoire compared with *K. rhizophila* M33 (Fig. 8B). Both genomes shared nitrogen-assimilation functions, including glutaminase, glutamine synthetase, glutamate synthase and glutamate dehydrogenase, while *G. arilaitensis* M33 was distinguished by a broader range of predicted aromatic-compound degradation pathways, including benzoate, catechol, protocatechuate, phenylacetate, 3-oxoadipate and salicylate 5-hydroxylase. Additional clusters involved in monoamine, tyramine/dopamine and xanthine metabolism further indicate greater metabolic versatility toward compounds potentially encountered in plant-associated environments. Its rCGC profile was also less strictly conserved across the genus, suggesting greater accessory functional diversity among *Glutamicibacter* genomes.

**Figure 8.**
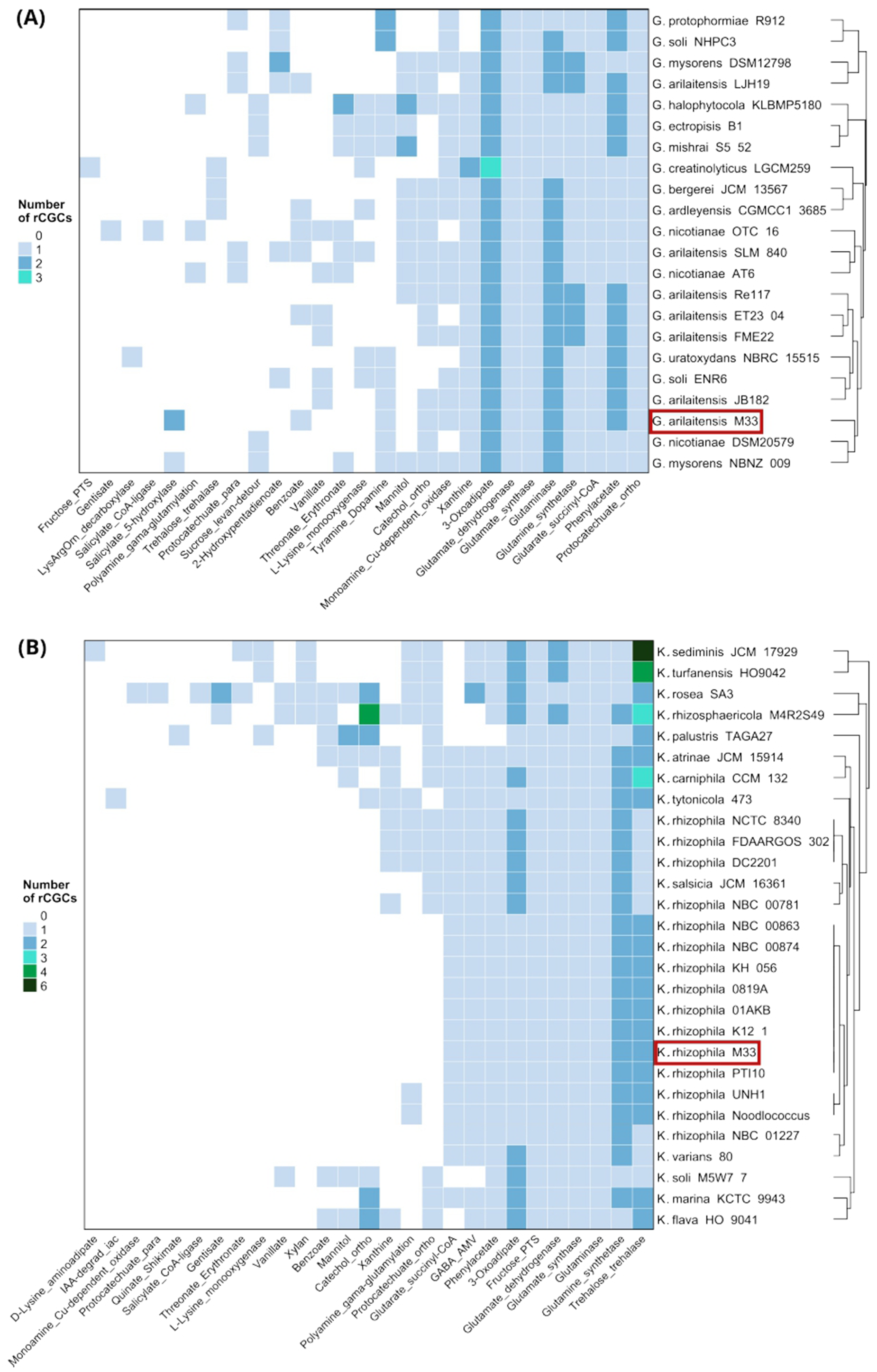
Distribution of rhizosphere-competence-related catabolic gene clusters (rCGCs) across *Kocuria* and *Glutamicibacter* genomes. Heatmaps show rhizoSMASH-predicted rCGCs in **(A)** 22 *Glutamicibacter* genomes and **(B)** 28 *Kocuria* genomes. Rows represent genomes and columns represent the 27 rCGC types detected within each genus. Genomes were clustered using Euclidean distances calculated from rCGC counts. Columns were ordered from the least prevalent rCGC types on the left to the most prevalent on the right, based on the number of genomes containing each cluster.

In contrast, *K. rhizophila* M33 clustered more closely with other *K. rhizophila* genomes, reflecting a highly conserved, species-specific rCGC profile. Its predicted functions were mainly associated with nitrogen and amino-acid metabolism, including glutamine synthetase, glutamate synthase, glutamate dehydrogenase, glutaminase and the GABA_AMV pathway. Additional clusters related to carbon acquisition and stress adaptation, such as fructose phosphotransferase system (PTS) and trehalose– trehalase systems, were also identified. Predicted 3-oxoadipate, phenylacetate and glutarate-succinyl-CoA pathways suggest a capacity to utilize a range of organic compounds, although its overall rCGC repertoire was less diverse than that of *G. arilaitensis* M33.

### 3.8 *K. rhizophila* M33 and *G. arilaitensis* M33 showed broad potential for plastic polymer degradation

Household waste disposal is not always accompanied by proper separation of organic waste; in particular, organic waste may be disposed of in conventional plastic bags rather than compostable bags, potentially affecting compost quality and possibly introducing microplastics into the final product.

We therefore investigated the potential of compost-isolated bacteria to metabolize plastic polymers. Using PlasticDB, the two M33 strains were screened for predicted plastic-degradation capabilities and compared with *Ideonella sakaiensis*, the first bacterium reported to degrade PET and a model organism for biological plastic recycling (Yoshida *et al*., 2016) (Figure 9, Table S14).

**Figure 9.**
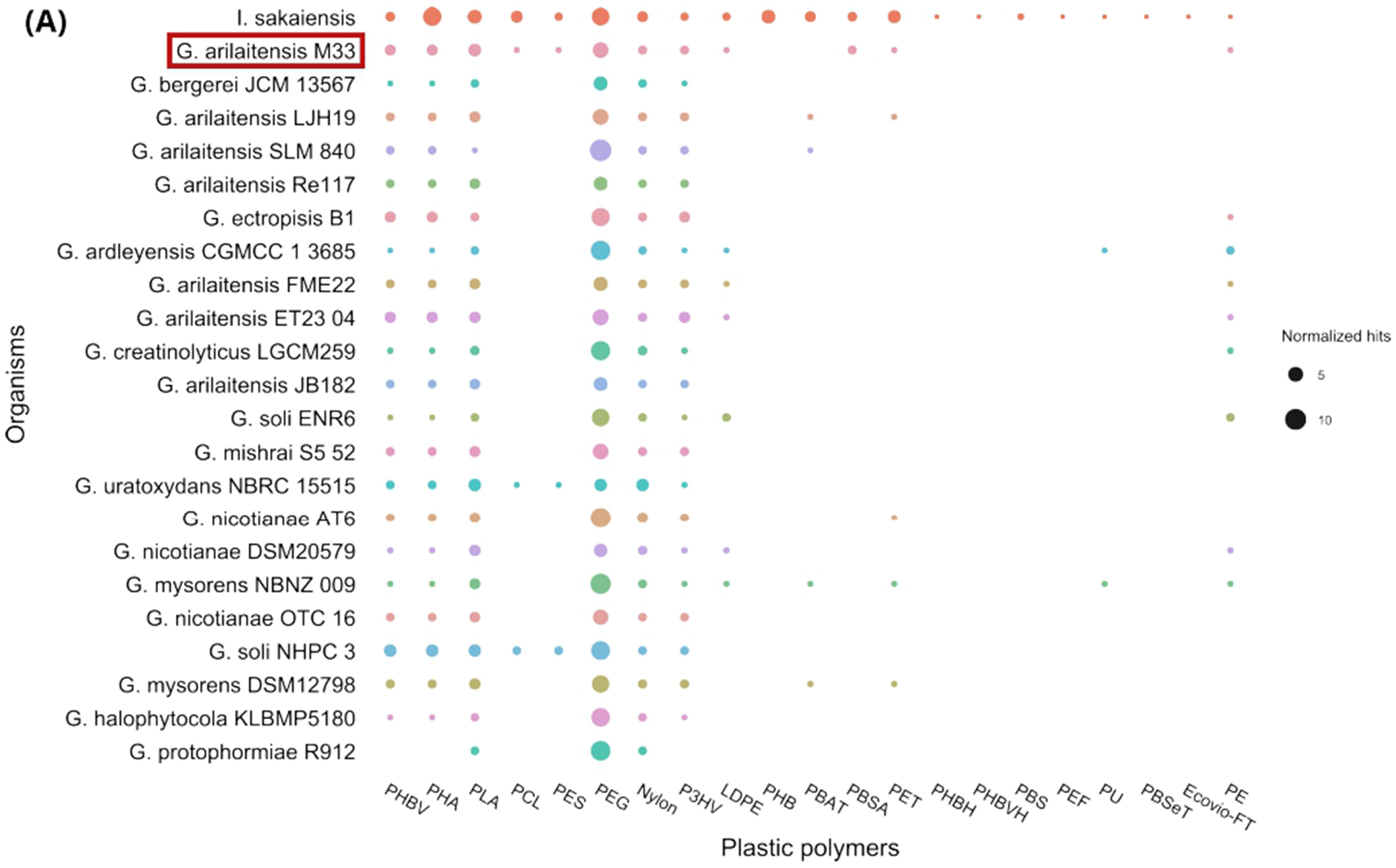
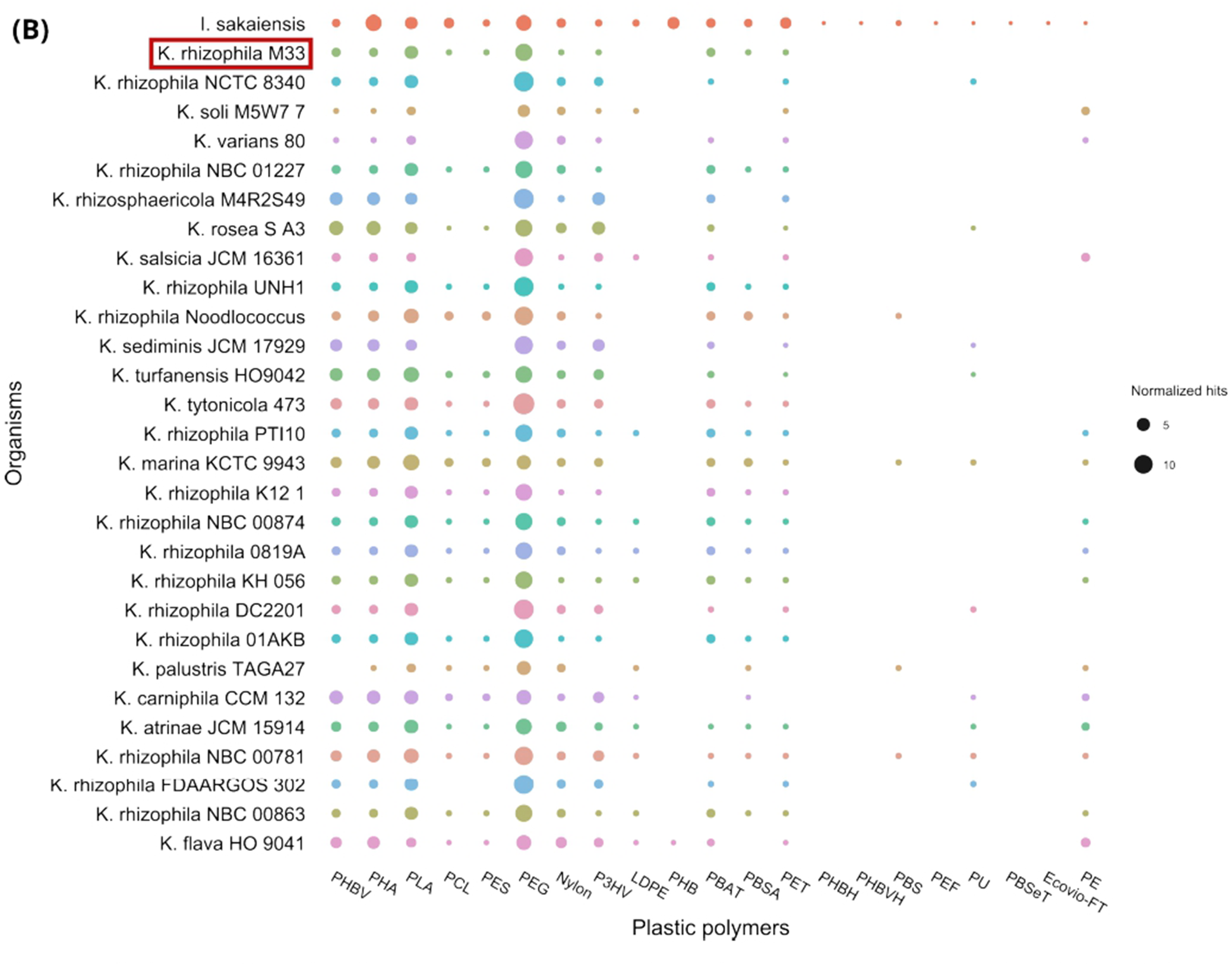
Distribution of putative plastic degradation associated protein homologues across *Kocuria* and *Glutamicibacter* genomes. Bubble plots summarize results obtained with PlasticDB for **(A)** 22 *Glutamicibacter* genomes and **(B)** 28 *Kocuria* genomes across 21 plastic polymer categories. *Ideonella sakaiensis* was included in both panels as a reference organism. Bubble size represents the number of PlasticDB hits normalized to genome protein content (total amino acids). Abbreviations: PHBV, poly(3-hydroxybutyrate-co-3-hydroxyvalerate); PHA, polyhydroxyalkanoate; PLA, polylactic acid; PCL, polycaprolactone; PES, polyethersulfone; PEG, polyethylene glycol; Nylon, nylon; P3HV, poly(3-hydroxyvalerate); LDPE, low-density polyethylene; PHB, polyhydroxybutyrate; PBAT, polybutylene adipate-co-terephthalate; PBSA, polybutylene succinate-co-adipate; PET, polyethylene terephthalate; PHBH, poly(3-hydroxybutyrate-co-3-hydroxyhexanoate); PHBVH, poly(3-hydroxybutyrate-co-3-hydroxyvalerate-co-3-hydroxyhexanoate); PBS, polybutylene succinate; PEF, polyethylene furanoate; PU, polyurethane; PBSeT, polybutylene sebacate-co-terephthalate; Ecovio FT, commercial blend of PLA and PBAT; PE, polyethylene.

Both strains showed a good potential to degrade different synthetic and biobased polymers. *G. arilaitensis* M33 (Fig. 9A) and *K. rhizophila* M33 (Fig. 9B) shared predicted degradation of PHBV, PHA, PLA, PCL, PES, PEG, nylon, P3HV, PBSA, and PET. Among these categories, PEG was associated with the greatest number of hits in both strains. LDPE and PE degradation was predicted only for *G. arilaitensis* M33 and was also observed in a small subset of closely related strains. Conversely, PBAT degradation was uniquely predicted for *K. rhizophila* M33 and was shared by nearly all its closest relatives, with the exception of three strains.

### 3.9 Genomic assessment of virulence and pathogenicity potential

To further assess the potential of *G. arilaitensis* M33 and *K. rhizophila* M33 as plant growth-promoting rhizobacteria, in particular their possible use as inoculants for plants intended for human consumption, virulence is an important aspect of safety assessment. Therefore, the genomes were further investigated for the presence of virulence-associated determinants and other features potentially related to pathogenicity.

The analysis conducted with VFDB produced matches to 23 VFDB reference sequences, represented by 38 hits in *G. arilaitensis* and 36 hits in *K. rhizophila* (Table S15 and S16). In *G. arilaitensis*, the strongest matches corresponded to the isocitrate lyase gene *icl*, the primary sigma-factor gene *sigA/rpoV*, and the nitrate-reductase genes *narH* and *narG*. Additional matches involved genes associated with stress responses and regulation, including *katG, relA, sigH, sigE, rpoS,* and *sodA*. The *K. rhizophila* genome showed its strongest matches to *icl* and *sigA/rpoV*. It also matched to the urease structural and accessory genes *ureA, ureC,* and *ureG*, together with genes involved in oxidative-stress resistance, nutrient acquisition, and transcriptional regulation, including *sodA, relA, sigH, mprA, ideR,* and *panD*.

Regarding the strongest hit, isocitrate lyase (Icl) is an enzyme of the glyoxylate cycle in *Mycobacterium tuberculosis* that contributes to fatty acid catabolism, intracellular replication, and persistence within the host (Muñoz-Elías & McKinney, 2005). Three *icl* hits were identified in each M33 strain. For this reason, to further assess their potential pathogenicity, the genomes were subsequently analysed using PathogenFinder2, which estimates the probability that a strain is pathogenic to humans. Both strains were predicted to be non-pathogenic to humans, with mean prediction scores of 0.125 ± 0.113 for *G. arilaitensis* and 0.274 ± 0.136 for *K. rhizophila* (Table S15 and S16).

### 3.10 Impact of *K. rhizophila* M33 inoculation on tomato plant growth and physiology

The previous biochemical and genomic characterizations supported the selection of *K. rhizophila* M33 as a candidate inoculant for compost enrichment and for assessing its potential effects on plant growth and health. Tomato plants were grown under different substrate treatments: soil, soil supplemented with compost, and soil supplemented with sterile compost, each with or without *K. rhizophila* M33 inoculation. Physiological responses were assessed 14 and 33 days after inoculation by measuring root and leaf fresh and dry biomass, photosynthetic efficiency, and leaf energy metabolism.

After 14 days, no visible differences in the physiological state of the plants were observed among the treatments, except for plants grown in sterile compost supplemented with *K. rhizophila*, which appeared slightly healthier (Figure S4A-F).

At 33 days, plant growth was strongly influenced by both the growing substrate and *K. rhizophila* inoculation (Figure S4G-L). Plants grown in soil showed the poorest development, whereas those grown in compost were substantially larger and greener. Inoculation with *K. rhizophila* further promoted growth in both soil and compost, resulting in taller plants, larger and darker green leaves, thicker stems, and more flowers. A similar positive effect was observed in sterile compost, where inoculated plants reached a development comparable to that of plants grown in compost with *K. rhizophila*.

Leaf dry weight measurements at 14 days post inoculation (dpi) were comparable among treatments (Figure 10A). At 33 dpi, leaf dry weight was higher in both compost-based substrates than in soil, with the highest and most variable values in sterile compost, though the differences were not statistically significant.

**Figure 10.**
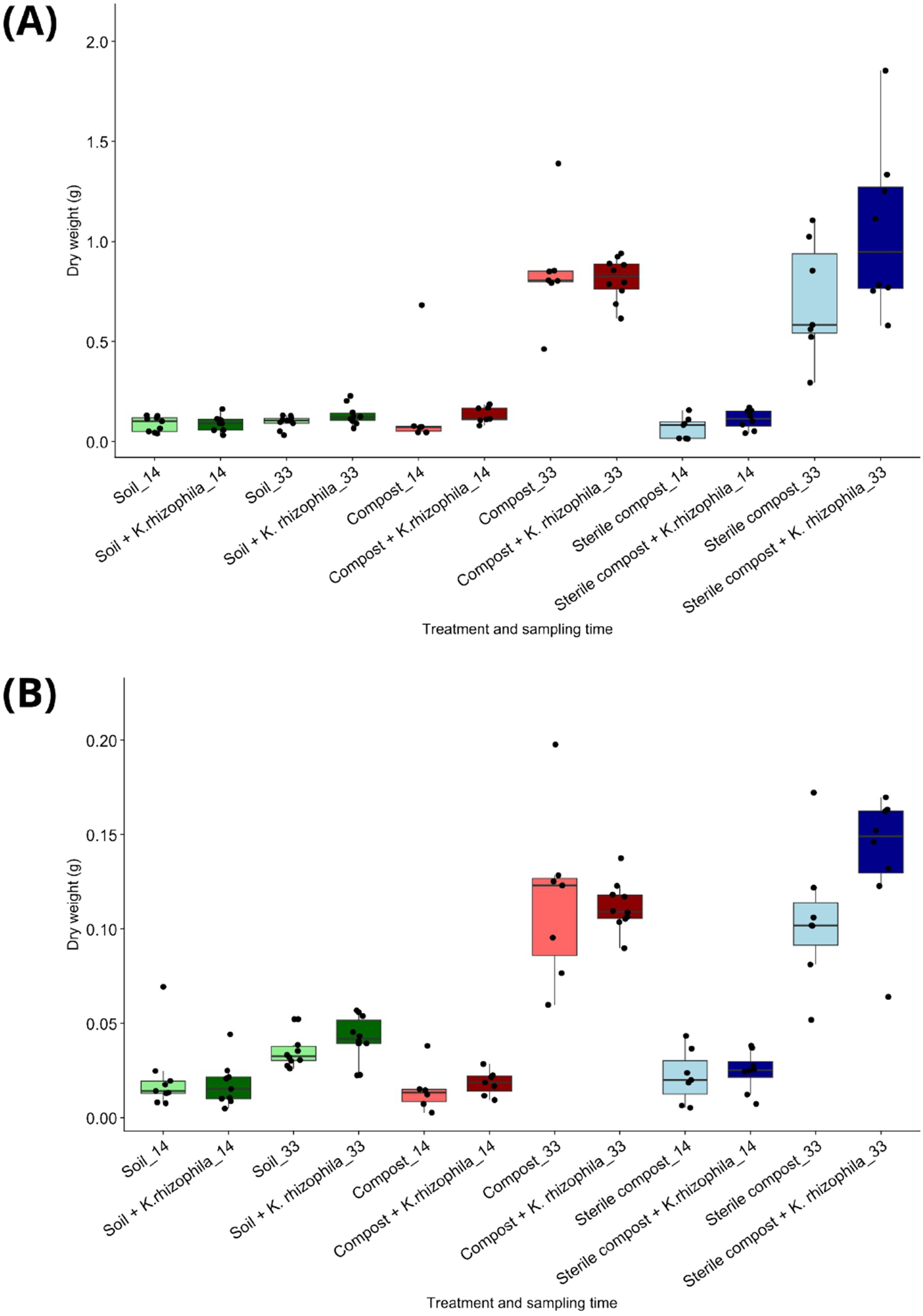
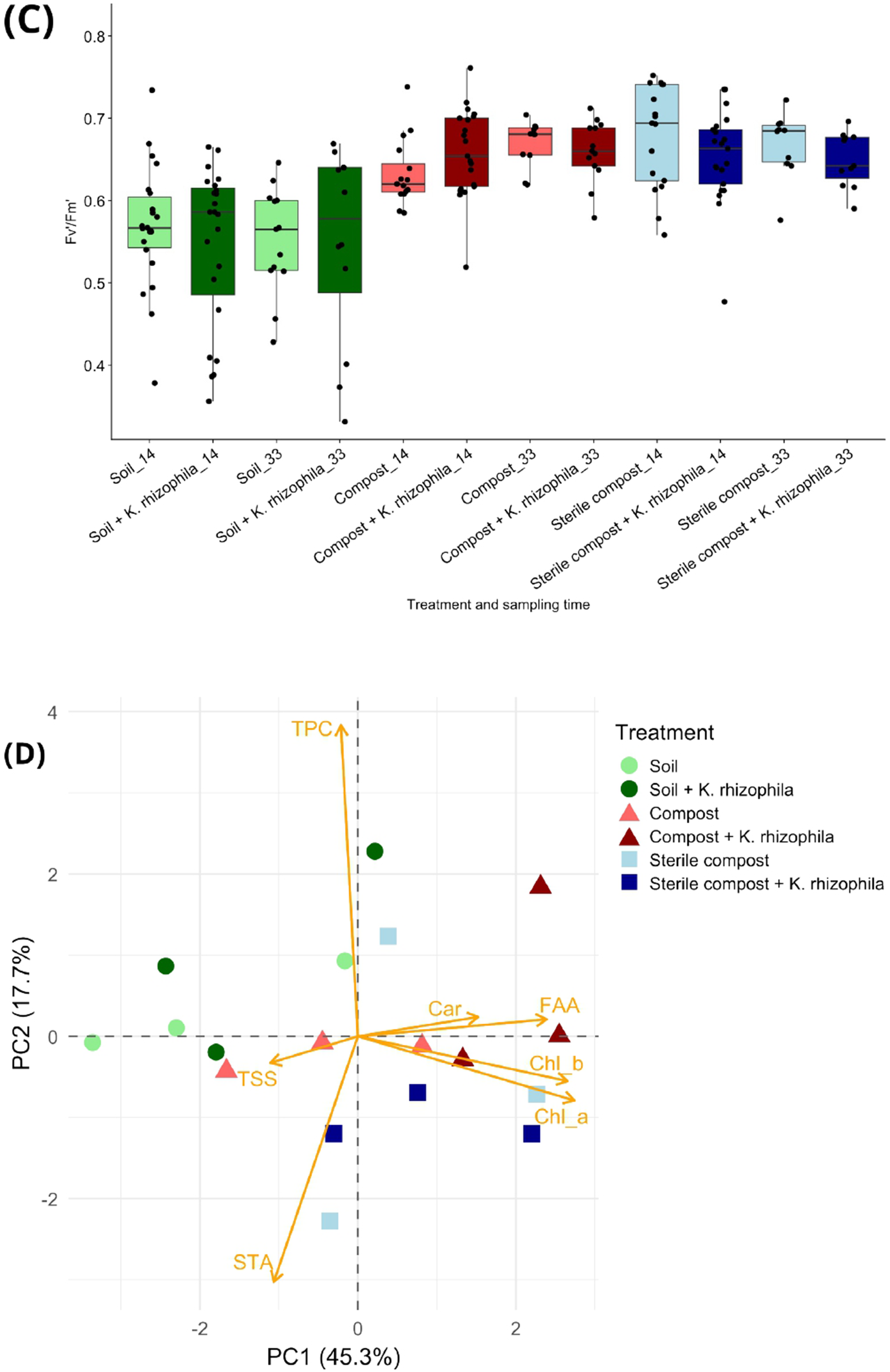
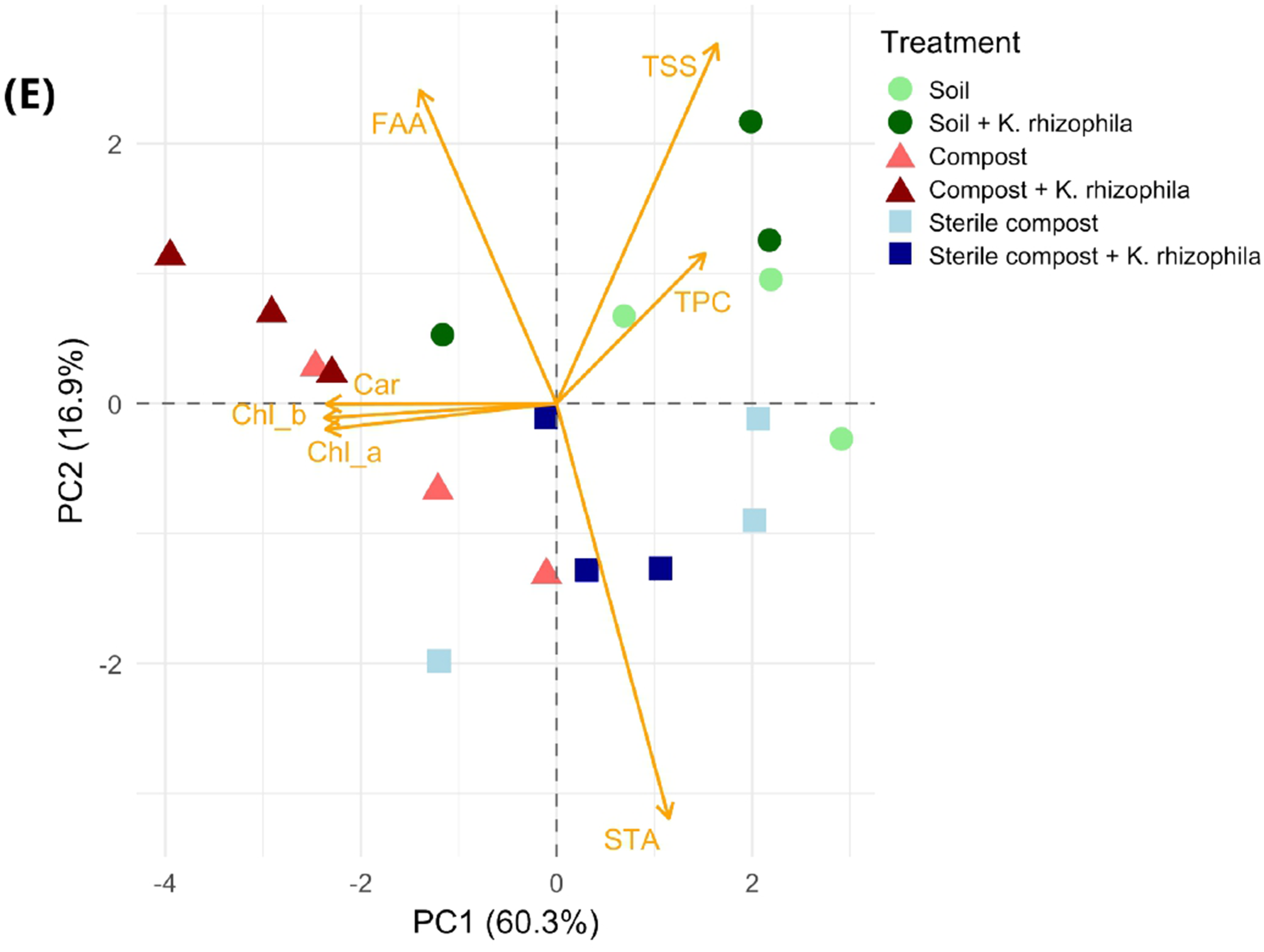
Effects of *K. rhizophila* M33 inoculation on tomato biomass, photosynthetic efficiency, and leaf metabolite profiles. Boxplots showing **(A)** leaf dry weight, **(B)** root dry weight and **(C)** photosynthetic efficiency (F_v_’/F_m_’) at 14 and 33 days post-inoculation (dpi). Boxes show the median and interquartile range, and points represent individual plants. **(D, E)** Principal component analysis of standardized leaf metabolite measurements at 14 and 33 dpi, respectively. Variables comprise free amino acids (FAA), carotenoids (Car), chlorophyll a and b (Chl_a and Chl_b), starch (STA), total phenolic compounds (TPC), and total soluble sugars (TSS).

Root dry weight showed a similar pattern (Fig. 10B), with limited differences at 14 dpi and higher values in compost-based substrates at 33 dpi. Inoculated plants in sterile compost M33 tended to have greater root dry weight than controls, but this difference was not significant. Overall, no significant inoculation effect on root dry weight was detected.

Fresh weight generally followed the dry-weight patterns (Figure S5). At 33 dpi, leaf fresh weight was higher in compost-based substrates than in soil, and inoculation significantly increased leaf fresh weight in sterile compost (Fig. S5A). No other significant effects of inoculation were observed for leaf or root fresh weight, although root fresh weight was higher in compost-based substrates (Fig. S5C).

Leaf water content was also calculated and reported for leaves (Fig. S5B) and roots (Fig. S5D), showing to be significantly higher in inoculated plants only in compost treatment at 33 dpi, while root water content was generally unaffected by inoculation.

To provide a more comprehensive assessment of plant health status, photosynthetic efficiency was also measured at 14 and 33 dpi (Fig. 10C). At both sampling times, values tended to be lower in soil-grown plants and higher in plants grown in compost and sterile compost, reflecting the nutritional availability of the different treatments. However, under the experimental conditions tested, inoculation did not produce a detectable change in photosynthetic efficiency, as the distribution of inoculated and non-inoculated plants largely overlapped within each substrate.

Principal component analysis was used to examine variation in the overall leaf metabolite profile (Fig. 10D-E). At 14 dpi, PC1 and PC2 explained 45.3% and 17.7% of the total variance, respectively, accounting together for 63.0% of metabolite variation (Fig. 10D). Samples showed substantial overlap, although some separation associated with substrate was apparent. After accounting for substrate, PERMANOVA detected no significant overall effect of *K. rhizophila* inoculation at this time point (p=0.128). The substrate × inoculation interaction was also not significant (p=0.210). At 33 dpi, the separation of metabolite profiles was more pronounced, with PC1 and PC2 explaining 60.3% and 16.9% of the variance, respectively, corresponding to 77.2% of the total variation (Fig. 10E). At this sampling time, PERMANOVA identified a significant effect of *K. rhizophila* inoculation after accounting for substrate (p=0.0285), which was associated with 9.9% of the overall variation in leaf metabolite profiles. This response was similar across the three substrates, as the substrate × inoculation interaction was not significant (Table S17).

## 4. Discussion

The application of biofertilizers represents an emerging strategy for more sustainable agriculture, but despite their rapid development, they remain a relatively small market segment, representing an estimated 0.7-1.4% of the global synthetic fertilizer market by value in 2022 (Dos Reis *et al*., 2024). Their effectiveness under field conditions remains highly variable and depends on soil and climatic conditions, the crop species and inoculant strain, formulation and application strategy, and interactions with resident soil microbial communities (Pirttilä *et al*., 2021; Ptaszek *et al*., 2023). Nevertheless, recent biotechnological advances and long-term field trials are increasingly supporting more targeted and integrated approaches to optimize their application to specific agricultural needs (Argentel-Martínez *et al*., 2025).

PGPR, which constitute an important component of microbial biofertilizers, are traditionally isolated from plant-associated environments, including the rhizosphere, rhizoplane, root endosphere, phyllosphere, and seeds. In contrast, the potential of organic fertilizers such as compost as a reservoir of beneficial, culturable bacteria remains largely unexplored. Although the positive contribution of compost-associated microbial communities to agricultural soil microbiota is well recognized, few studies have isolated and characterized individual compost-derived bacteria as potential bioinoculants. Recent work has reported beneficial bacteria from plant-based and/or homemade compost (Suárez-Estrella *et al*., 2019; Kim *et al*., 2021; Vásquez & Millones, 2023), whereas our study extends this approach to five batches of commercially produced industrial compost derived from municipal organic waste.

In this work, we isolated and identified six culturable bacterial strains, three of which belonged to the genus *Bacillus*. This finding is consistent with the high relative abundance of the Bacillaceae family observed in the compost bacterial community (unpublished data). All isolated strains were Gram-positive, which reflects their adaptation to the selective conditions of the composting process: composting involves high temperatures, reaching up to 70 °C, and therefore favours bacteria capable of forming heat-resistant endospores, such as *Bacillus*. Other isolated genera, including *Kocuria*, *Glutamicibacter*, and *Microbacterium*, belong to the phylum Actinomycetota and they can grow in compost under both mesophilic and thermophilic conditions, at temperatures of up to 50-60 °C and under neutral to alkaline pH (Aguilar-Paredes *et al*., 2023). Furthermore, the compost used in this study exhibited relatively high salinity, and the isolated species have been reported to tolerate saline conditions and, in some cases, to contribute to the alleviation of salt stress in plants (Afridi *et al*., 2021; Hidri *et al*., 2023; Valencia-Marin *et al*., 2024).

All strains except *Bacillus licheniformis* exhibited two or three of the plant growth-promoting traits tested. *Bacillus* species are widely recognized for their beneficial effects on plants, including protection against both biotic and abiotic stresses (Tsotetsi *et al*., 2022). In this study, the biochemical assays focused on auxin production and nutrient solubilization; therefore, the lack of detectable PGP traits in *B. licheniformis* does not necessarily indicate an absence of beneficial activity. Indeed, this strain may possess other functions not assessed here, such as biocontrol activity, which has been reported for other plant-associated strains of *B. licheniformis* (Nigris *et al*., 2018; Ji *et al*., 2023).

The biochemical profiles of *K. rhizophila* and *G. arilaitensis* supported whole-genome sequencing of the two strains, which enabled the robust taxonomic placement and provided a basis for investigating their genomic variability and potential adaptive traits.

Specifically, *G. arilaitensis* M33 was most closely related to ET23-04, a metagenome-assembled genome from a marine biofilm (NCBI, 2026), highlighting that genomic relatedness does not necessarily reflect ecological origin. Our *G. arilaitensis* M33 strain can instead be more directly compared with *G. arilaitensis* LJH19, which was isolated from a Himalayan night-soil composting system and exhibits several plant growth-promoting traits similar to those investigated here (Borker *et al*., 2021). The phylogenomic proximity of *K. rhizophila* M33 to soil-derived strains NBC_00863 and NBC_00874 suggests a possible association with terrestrial, nutrient-rich environments; however, the distinct position of M33 strain within the clade reflects strain-level divergence and possible adaptation to compost-specific conditions.

Most predicted genes in both M33 genomes were assigned to orthogroups shared with related genomes, indicating that their overall gene repertoires are largely conserved within their respective phylogenetic groups. Among annotated genes, the representation of amino acid and carbohydrate transport and metabolism is compatible with the ability to use a range of nutrients, which could be advantageous in a chemically heterogeneous environment such as compost.

Moreover, both isolates contained a comparatively small set of candidate M33-specific genes. Only a proportion of the candidate M33-specific genes could be assigned to COG functional categories, limiting conclusions about their biological significance, requiring further validation.

Overall, this genomic characterization confirmed the presence of genes associated with several of the PGP traits detected through biochemical assays. Interestingly, *G. arilaitensis* M33 showed a predicted genetic potential for phosphate solubilization, although no corresponding activity was detected phenotypically. One of the close relatives of this strain, *G. arilaitensis* LJH19, has been instead reported to exhibit phosphate-solubilizing activity (Borker *et al*., 2021), highlighting the importance of strain-specific validation of genomic predictions.

Further analysis of rCGCs revealed both shared and strain-specific functional patterns between the two M33 isolates. This information may be relevant for the design of bacterial consortia, as strains with overlapping catabolic capabilities may possess a competitive advantage during co-cultivation and plant colonization (Chen & Liu, 2024). Conversely, complementary metabolic profiles could favour coexistence and functional cooperation within a consortium.

Beyond their potential roles in plant-associated microbial communities, the metabolic versatility of these compost-derived strains may also be relevant to the degradation of other organic and synthetic compounds present in composting environments. In Italy, an estimated 7.1% of the organic fraction of municipal waste sent to composting facilities in 2022 contained non-compostable or unsuitable materials, including plastic waste (Centro Studi CIC, 2024). Given this issue, we used PlasticDB to investigate the potential of our compost-derived bacteria to degrade plastic polymers. Compared with the well-characterized plastic-degrading bacterium *Ideonella sakaiensis*, both M33 strains showed predicted capabilities for approximately half as many plastic polymer categories. These results suggest that, although less versatile than *I. sakaiensis*, the two compost-derived strains may possess a relatively broad plastic-degradation potential.

Interestingly, plastic-degradation capabilities have also been reported in closely related taxa. Two bacteria isolated from landfill soils, including *Glutamicibacter mysorens* and a closely related *Arthrobacter* strain, have demonstrated plastic-degrading activity both experimentally and through genomic analyses (Velo *et al*., 2025; Sherigar *et al*., 2026). Similarly, a landfill-derived *K. rhizophila* strain was reported to degrade polyurethane (Muñoz-Martí *et al*., 2025), while *Kocuria palustris* has been shown to degrade low-density polyethylene (Harshvardhan & Jha, 2013). Together, these findings support the potential of *Kocuria* and *Glutamicibacter* species as promising candidates for further investigation of plastic biodegradation, while experimental validation will be required to confirm the predicted activities in M33. To conclude the genetic characterization, and given the potential application of the M33 strains as crop bioinoculants, their potential risk to human health was assessed.

Both *G. arilaitensis* and *K. rhizophila* are predominantly environmental bacteria and are not generally regarded as pathogenic, although rare opportunistic infections have been reported for some members of both genera, particularly in immunocompromised individuals. No established pathogenic role has been reported for *G. arilaitensis*, and compost-derived *G. arilaitensis* LJH19 has previously shown no haemolytic activity and has been proposed as a potential safe plant growth-promoting inoculant. Similarly, *K. rhizophila* has only rarely been associated with human infections (Ziogou *et al*., 2023). In the present study, neither M33 strain was predicted to be pathogenic based on genomic analyses, supporting a favourable preliminary safety profile for both strains.

*In vivo* testing of bioinoculants is an important approach for gaining direct insights into their effects on plant growth and physiology. Controlled growth-chamber experiments provide a valuable environment to assess plant responses to bacterial inoculation and optimize inoculation strategies before field application. In this study, we investigated the effects of *K. rhizophila* M33 on tomato plant physiology under different substrate conditions, including soil, soil supplemented with compost, and soil supplemented with sterile compost. The use of compost enriched with bacteria originally isolated from the same matrix may represent a promising strategy, as these strains are potentially pre-adapted to the physicochemical conditions of the compost, which could favour their establishment and contribute to more efficient compost utilization.

Although the observed responses were influenced more strongly by the substrate than by bacterial inoculation, *K. rhizophila* M33 treatment was associated with a trend towards increased biomass, particularly in plants grown in sterile compost. This effect may be attributable to reduced competition from the resident microbial community, which could have enhanced the activity or persistence of the inoculated strain. In contrast, the effects on leaf metabolism were less pronounced, suggesting a moderate metabolic modulation rather than extensive reprogramming. The limited metabolic response may also indicate that the effects of *K. rhizophila* M33 were not strongly reflected in leaf metabolism under the conditions tested. These findings are consistent with recent studies showing that sugars, amino acids, and energy regulators involved in tomato growth were strongly modulated upon *K. rhizophila* inoculation in pot experiments (Mauceri *et al*., 2024). Similarly, *K. rhizophila* isolated from an arid grassland environment improved tomato plant productivity and fruit yield under both irrigated and water-deficit conditions (Almalkawi *et al*., 2025). Although, to our knowledge, no previous studies have investigated the effects of *G. arilaitensis* inoculation on tomato plants, the genomic characterization performed in this study revealed a range of potentially beneficial traits that support its consideration as a promising candidate for future in vivo evaluation.

Overall, this study highlights industrial biowaste-derived compost as a promising source of bacteria with potential applications in sustainable agriculture. The combination of genomic, phenotypic, and in vivo approaches provides a basis for further developing compost-derived strains as bioinoculants. Together, these findings support a circular strategy in which bacteria recovered from compost are used to enhance the biological value of the same organic amendment. An interesting next step would be to explore bacterial consortia combining the two M33 strains, to investigate their overall effectiveness in tomato plant performance. Moreover, further experiments under field-relevant conditions will be important to assess their practical potential.

## Authors’ contributions

Conceptualization, SN and RC; methodology, RC and SN; formal analysis, RC, SRS, NdB and CLA; investigation, NdB, RC, GG and CLA; resources, SN, BB, SRS, TB and WZ; data curation, RC and SRS; supervision, SN and BB; funding acquisition, SN and BB; writing - original draft preparation, RC; writing - review and editing, SN, RC and BB. All authors reviewed and approved the final version of the manuscript.

## Acknowledgements

We thank Leonardo Greggio, MSc, for his contribution to the growth curve experiments; Prof. Marco Giovannetti (University of Turin, Italy) and Prof. Dr. Gabriele Berg (Graz University of Technology, Graz, Austria), doctoral advisors of RC, for their valuable suggestions.

## Data availability statement

The sequence data generated during this study have been deposited in the NCBI GenBank database under the BioProject PRJNA1545693.

The supplementary tables are available on Zenodo

(https://zenodo.org/records/23536792)

