## Supplementary Figures for "Genomic and functional characterization of compost-derived bacteria identifies a candidate bioinoculant that influences tomato physiology and leaf metabolism"

### Supplementary Tables

The supplementary tables are available on Zenodo (*https://zenodo.org/records/23536792*)

**Table S1**. Physicochemical properties of compost batches M33, M38, M39, M40, M41 and M42 assessed by S.E.S.A. S.p.A.

**Table S2**. Compost humidity calculated by S.E.S.A. S.p.A.

**Table S3**. Indole-3-acetic acid standard-curve measurements used for IAA quantification.

**Table S4**. Soil physicochemical properties assessed by S.E.S.A. S.p.A.

**Table S5**. Compost CFU analysis results.

**Table S6**. Taxonomic identification of bacterial isolates based on nucleotide-sequence BLAST analysis.

**Table S7**. Growth kinetics and viable cell counts of the selected bacterial isolates during 28 h of cultivation.

**Table S8**. Quantification of indole-3-acetic acid production by the selected bacterial strains.

**Table S9**. OrthoFinder summary statistics for *Glutamicibacter arilaitensis* M33 and 21 reference genomes.

**Table S10**. OrthoFinder summary statistics for *Kocuria rhizophila* M33 and 27 reference genomes.

**Table S11**. BLASTn results for the putative plasmid contig identified in the *Glutamicibacter arilaitensis* M33 genome assembly.

**Table S12**. Putative antimicrobial resistance associated matches detected in the putative plasmid of *Glutamicibacter arilaitensis* M33 using CARD RGI.

**Table S13**. Biosynthetic gene clusters predicted using antiSMASH in the two M33 strains.

**Table S14**. Plastic-degradation-associated protein hits identified using PlasticDB in the *Kocuria rhizophila* M33 and *Glutamicibacter arilaitensis* M33 comparative genome datasets.

**Table S15**. Virulence-associated features and pathogenicity predictions for the *Glutamicibacter arilaitensis* M33 using the Virulence Factor Database (VFDB) and PathogenFinder2.

**Table S16**. Virulence-associated features and pathogenicity predictions for the *Kocuria rhizophila* M33 using the Virulence Factor Database (VFDB) and PathogenFinder2.

**Table S17**. Principal component analysis and PERMANOVA of leaf biochemical profiles at 14 and 33 days post-inoculation.

### Supplementary Figures

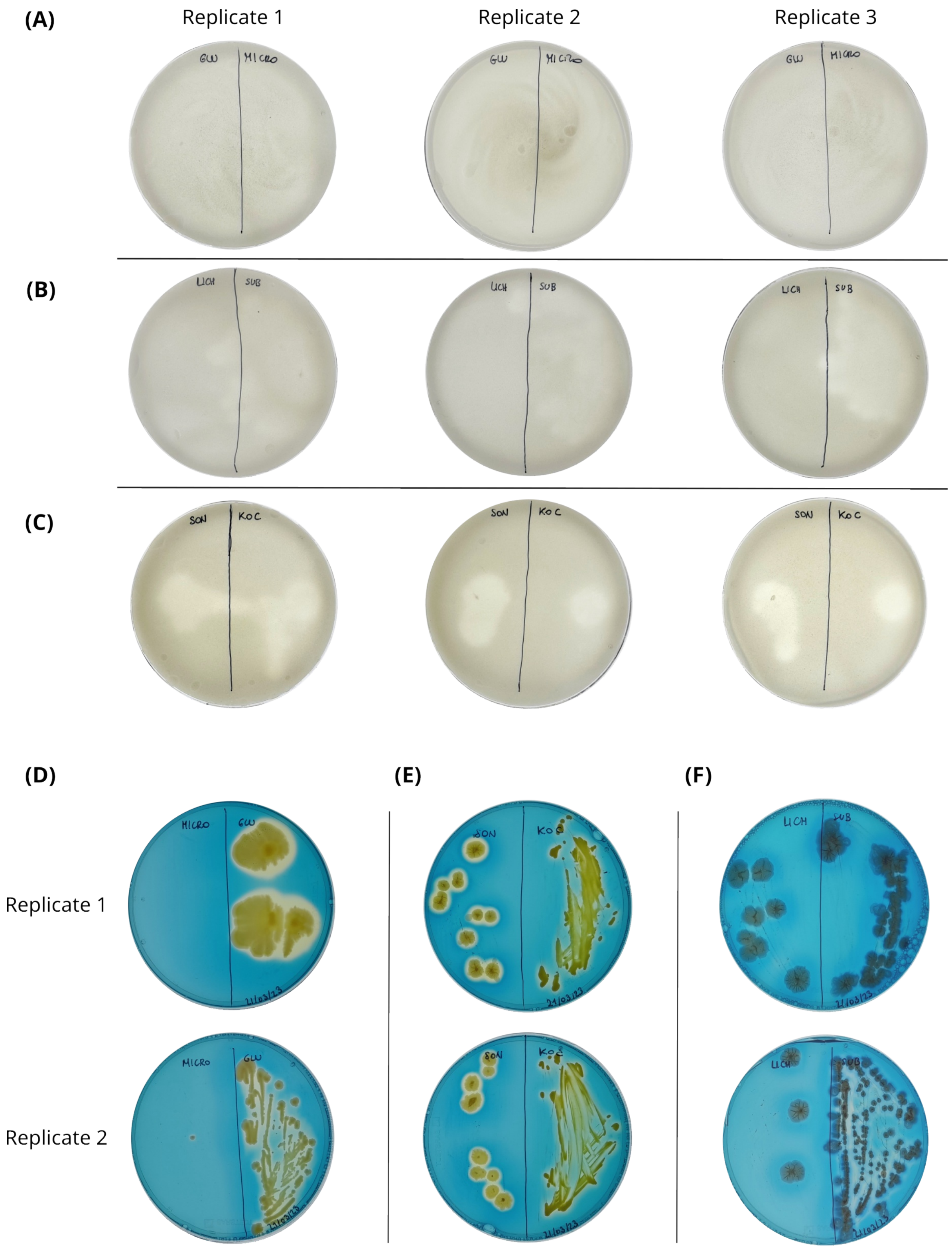


**Figure S1. Qualitative assessment of phosphate solubilization and siderophore production by the selected bacterial isolates.** Upper panels show phosphate solubilization on Pikovskaya agar: **(A)** *Glutamicibacter* sp. (left) and *M. suwonense* (right), **(B)** *B. licheniformis* (left) and *B. subtilis* (right), and **(C)** *B. sonorensis* (left) and *K. rhizophila*. Lower panels show siderophore production on chrome azurol S (CAS) agar: **(D)** *M. suwonense* (left) and *Glutamicibacter* sp. (right), **(E)** *B. sonorensis* (left) and *K. rhizophila* (right), and **(F)** *B. licheniformis* (left) and *B. subtilis* (right). Zones of clarification on Pikovskaya agar indicate phosphate solubilization, whereas a color change from blue to yellow-orange on CAS agar indicates siderophore production. Three and two replicate plates are shown for the phosphate-solubilization and siderophore-production assays, respectively.

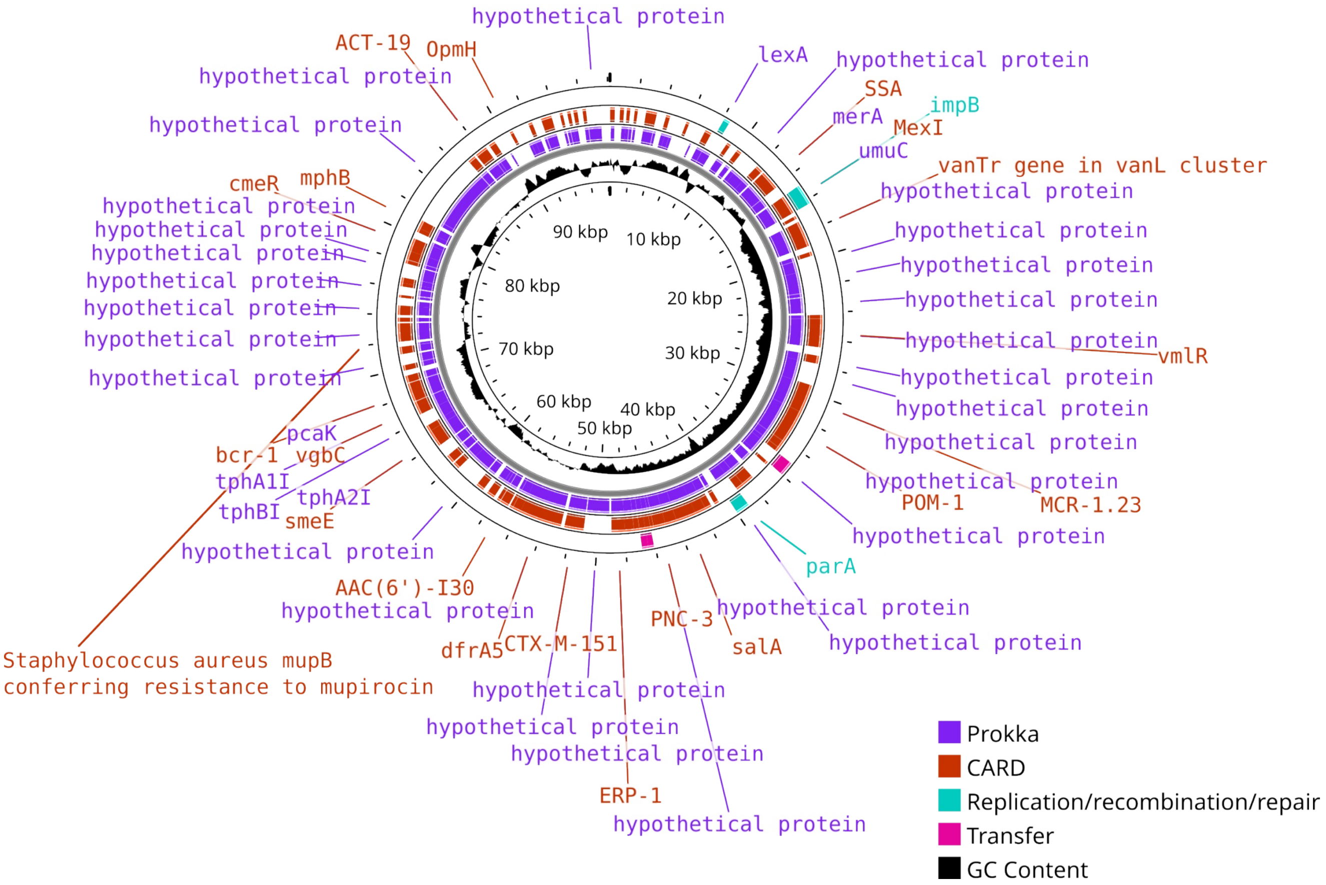


**Figure S2. Circular map of the putative plasmid contig identified in the *G. arilaitensis* M33 genome assembly.** Circular genome map visualized with Proksee showing coding sequences annotated with Prokka, antimicrobial resistance-associated determinants, and mobile genetic element-associated loci. Arrows indicate the positions and orientations of coding sequences, while the inner track represents local GC content.

**(A)**

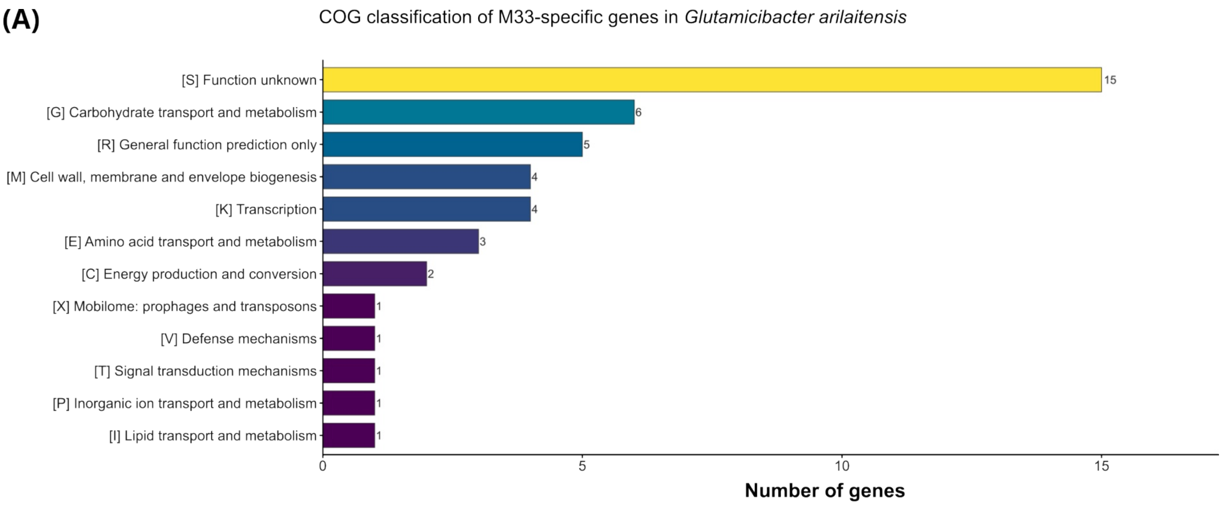


**(B)**

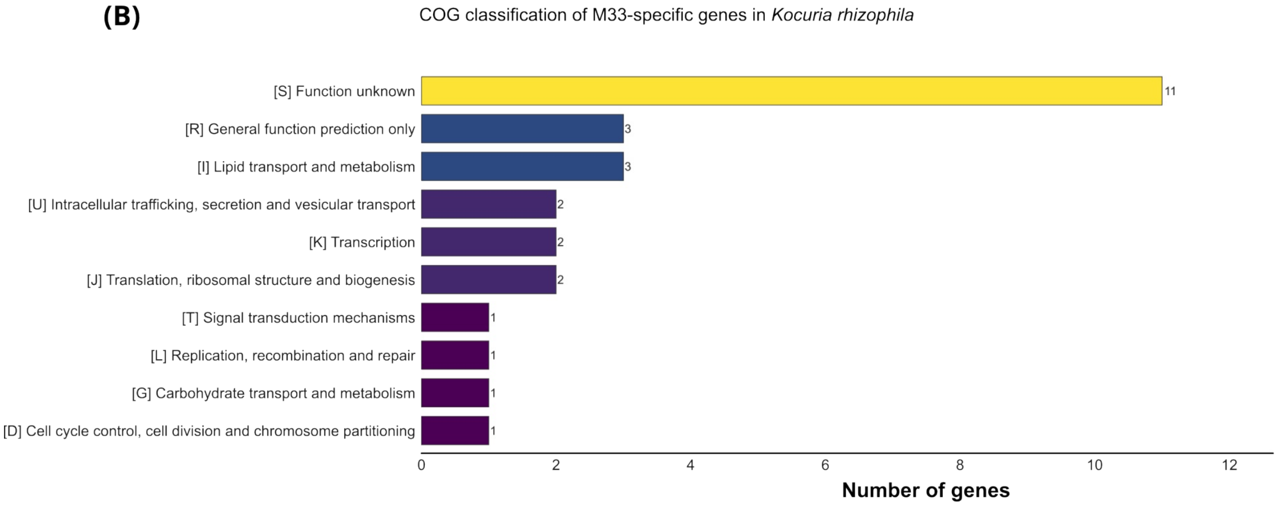


**Figure S3. COG classification of M33-specific genes identified by comparative genomic analysis.** Candidate M33-specific genes were defined as genes belonging to OrthoFinder orthogroups present in M33 and absent from the ten closest genomes, together with genes not assigned to an orthogroup. **(A)** *G. arilaitensis* M33. **(B)** *K. rhizophila* M33. Genes were functionally annotated using eggNOG-mapper and grouped according to their COG categories. Bars indicate the number of genes assigned to each category. Genes associated with multiple COG categories were counted once in each corresponding category.

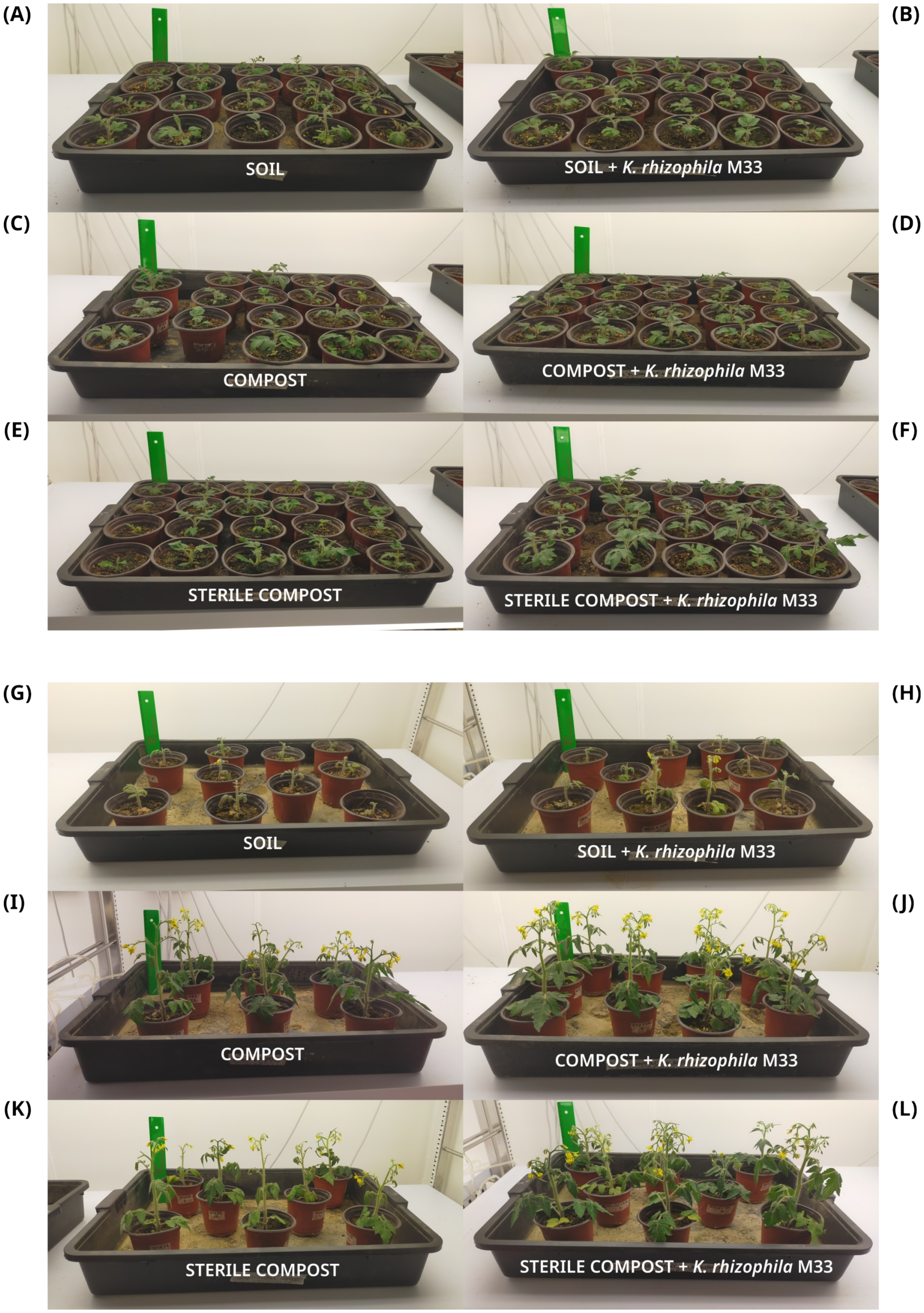


**Figure S4. Growth of tomato plants in different substrates with or without *K. rhizophila* M33 inoculation.** Plants are shown at 14 days post-inoculation (dpi) in **(A)** soil, **(B)** soil with *K. rhizophila* M33, **(C)** compost, **(D)** compost with *K. rhizophila* M33, **(E)** sterile compost, and **(F)** sterile compost with *K. rhizophila* M33. The corresponding treatments at 33 dpi are shown in panels **(G–L)**, respectively.

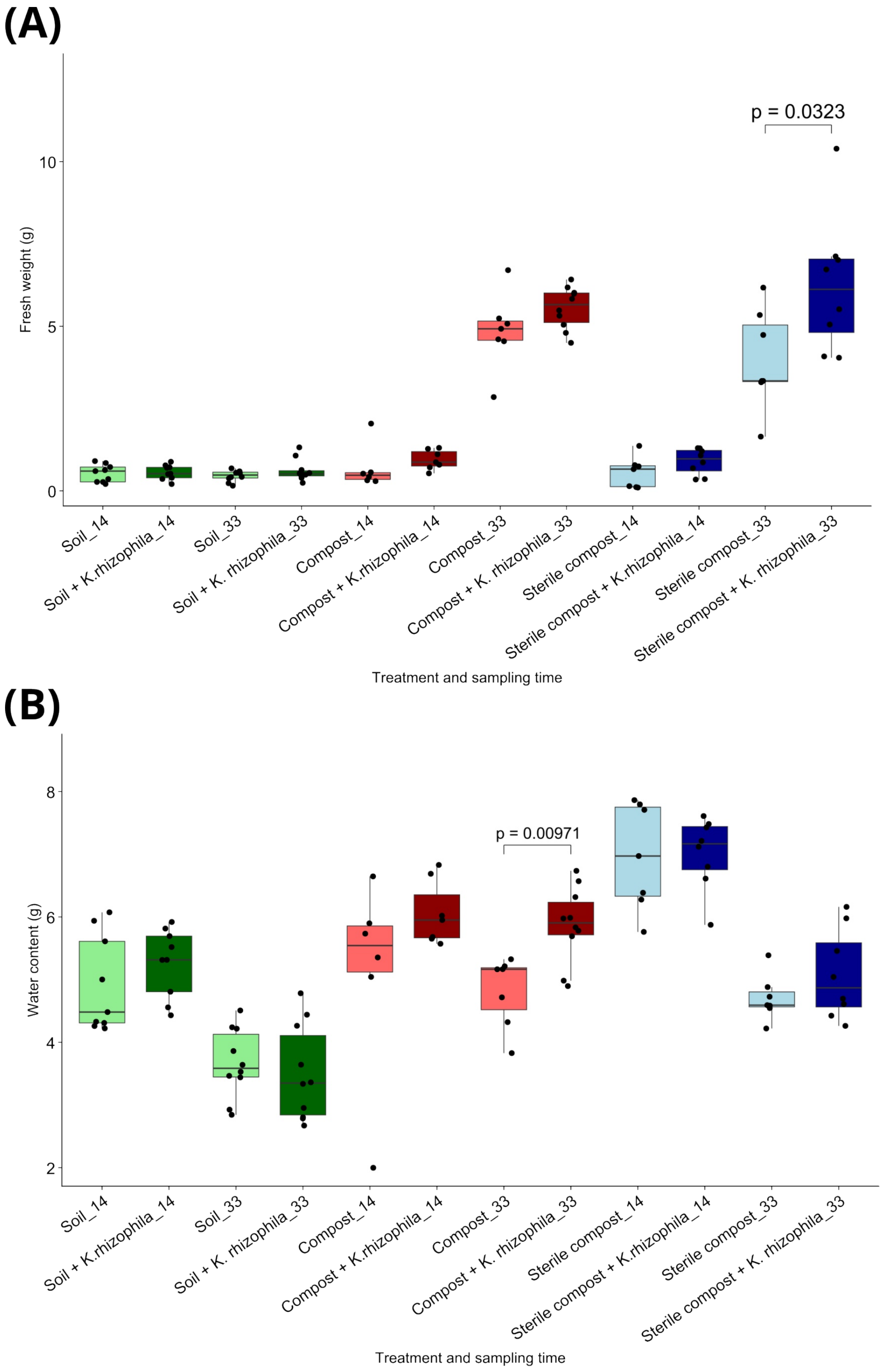


**Figure S5. Effects of *K. rhizophila* M33 inoculation on tomato fresh biomass and water content.** Boxplots showing **(A)** leaf fresh weight, **(B)** leaf gravimetric water content at 14 and 33 days post-inoculation (dpi) under the different soil conditions. Statistical differences between uninoculated and inoculated plants within each substrate and sampling time were assessed using two-sided Wilcoxon rank-sum tests. Differences were considered statistically significant at $p < 0.05$.

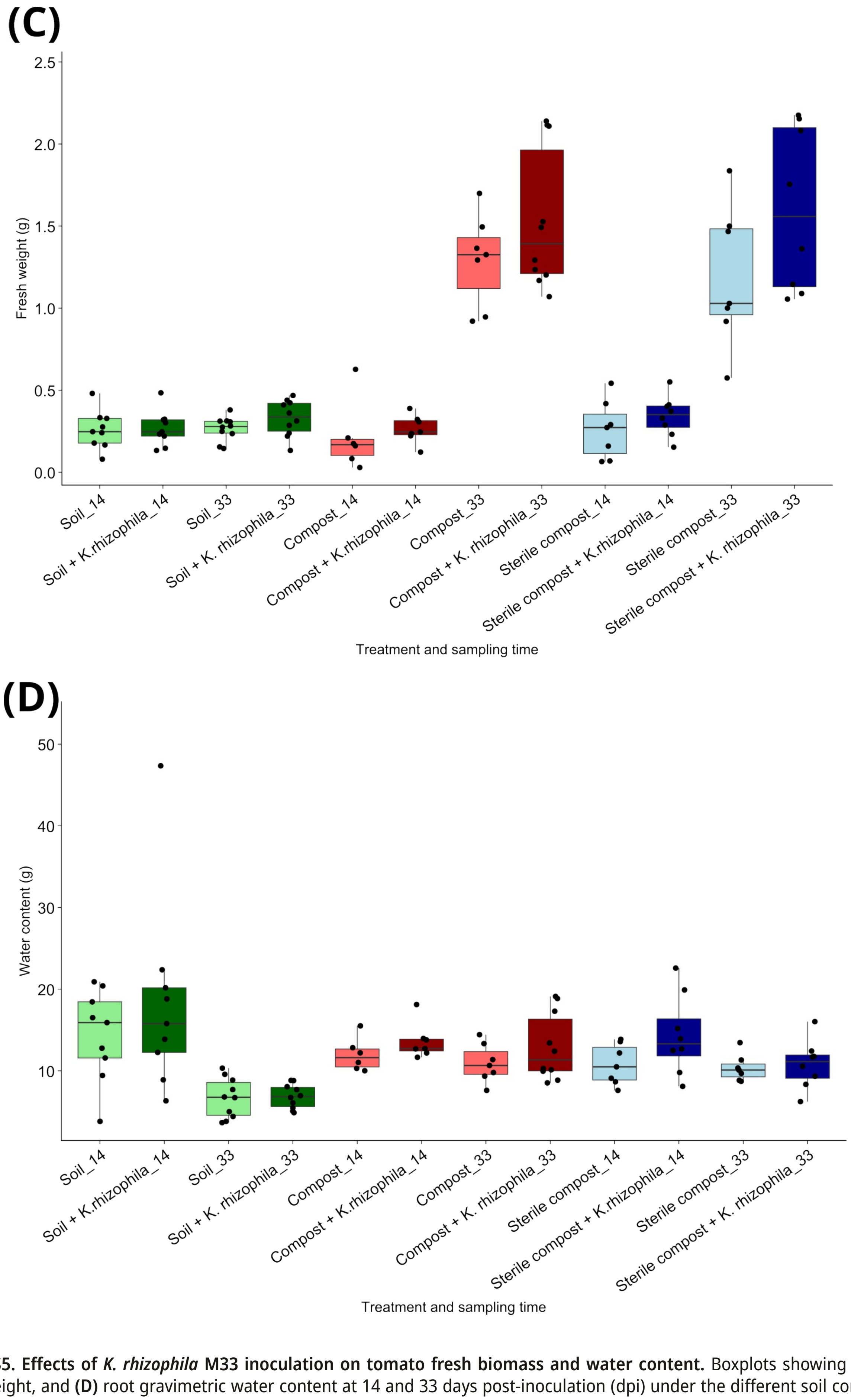


**Figure S5. Effects of *K. rhizophila* M33 inoculation on tomato fresh biomass and water content.** Boxplots showing **(C)** root fresh weight, and **(D)** root gravimetric water content at 14 and 33 days post-inoculation (dpi) under the different soil conditions. Statistical differences between uninoculated and inoculated plants within each substrate and sampling time were assessed using two-sided Wilcoxon rank-sum tests. Differences were considered statistically significant at $p < 0.05$.
